# B1 SINE-binding ZFP266 impedes reprogramming through suppression of chromatin opening mediated by pioneering factors

**DOI:** 10.1101/2022.01.04.474927

**Authors:** Daniel F Kaemena, Masahito Yoshihara, James Ashmore, Meryam Beniazza, Suling Zhao, Mårten Bertenstam, Victor Olariu, Shintaro Katayama, Keisuke Okita, Simon R Tomlinson, Kosuke Yusa, Keisuke Kaji

## Abstract

Induced pluripotent stem cell reprogramming is inherently inefficient and understanding the molecular mechanisms underlying this inefficiency holds the key to successfully control cellular identity. Here, we report 16 novel reprogramming roadblock genes identified by CRISPR/Cas9-mediated genome-wide knockout (KO) screening. Of these, depletion of the predicted KRAB zinc finger protein (KRAB-ZFP) *Zfp266* strongly and consistently enhanced iPSC generation in several iPSC reprogramming settings, emerging as the most robust roadblock. Further analyses revealed that ZFP266 binds Short Interspersed Nuclear Elements (SINEs) adjacent to binding sites of pioneering factors, OCT4 (POU5F1), SOX2 and KLF4, and impedes chromatin opening. Replacing the KRAB co-suppressor with a co-activator domain converted ZFP266 from a reprogramming inhibitor to a potent reprogramming facilitator. This work proposes SINE-KRAB-ZFP interaction to be a critical regulator of chromatin accessibility at enhancers for efficient cellular identity changes and also serves as a resource to further illuminate molecular mechanisms hindering reprogramming.

## Introduction

The reprogramming of somatic cells into iPSCs via the overexpression of *Oct4 (Pou5f1), Sox2, Klf4* and *c-Myc* (OSKM) has provided an important tool for medical research and cell therapies^1^. Equally importantly, the generation of fully functional iPSCs that are indistinguishable from ESCs from somatic cells has demonstrated that cellular identity can be completely converted from one type to another by overexpression of master transcription factors. This has provided a model system to understand how to control cellular identity. Inhibition of *Trp53* and *Cdkn1a* (*p21*) revealed OSKM-induced apoptosis and senescence as a major roadblock of iPSC generation^2–8^. Knockdown of *Dot1l* and *Suv39h1* has demonstrated H3K79me and H3K9me3 as critical epigenetic modifications that impede this cell conversion^9,10^. Thus, identifying genes that act against successful reprogramming provides the foundation to understand critical molecular mechanisms involved in pluripotency induction.

Transposable elements (TEs), which constitute approximately 40% of mouse and human genomes, take part in gene expression regulation as cis-regulatory elements or non-coding RNAs^11^. Long terminal repeat (LTR) retrotransposons, long interspersed elements (LINEs), and SINEs are the three major classes of human/mouse TEs and the functional importance of the first two groups in pluripotent cells has been described^12,13^. Knockdown of the long interspersed element 1 (LINE1) inhibits mouse ESC self-renewal and induces transition to a 2C state^12^. KLF4 activates transcription of LTR retrotransposon human endogenous retrovirus subfamily H (HERVH) during reprogramming, and the down-regulation of which is critical for exit from the pluripotent state of human iPSCs^13^. Chromatin accessibility of SINEs, which constitute ~25% of TEs, is particularly high in mouse pre-implantation embryos and ESCs^14^, but the functional importance of this has not been demonstrated yet. Krüppel-associated box (KRAB) zinc-finger proteins (ZFPs) form the largest TF family in mouse and human genomes with over 300 members^15^. They have evolved to supress expression and transposition of rapidly mutating TEs, with about two thirds of human KRAB-ZFPs estimated to bind to TEs^16^. Thus, some of KRAB-ZFPs might be involved in the regulation of the above mentioned pluripotency-associated LINE1 and HERVK expression. Binding of KRAB-ZFPs on TEs can also regulate the expression of nearby genes^17^. Knockout of the KRAB-ZFP cluster in chromosome 2 or chromosome 4, which contains 40 or 21 KRAB-ZFPs, respectively, in mouse ESCs preferentially up-regulated genes near specific classes of LTR retrotransposons and LINEs^18^. Overexpression of ZNF611 in human ESCs down-regulated genes near primate specific SINE-VNTR-Alu (SVA) retrotransposons^19^. Nevertheless, only a small number of KRAB-ZFPs that predominantly bind SINEs have been reported^16,18^, and the importance of KRAB-ZFP/SINE interaction for gene expression regulation is not well understood.

Here, we report an unbiased genome-wide CRISPR KO screen with a library containing 90,230 sgRNAs targeting 18,424 protein coding genes. This screen identified 16 genes as novel reprogramming roadblocks, as well as 8 previously reported roadblock genes. Of those, KO of the previously uncharacterised KRAB-ZFP gene *Zfp266* accelerated the kinetics of reprogramming and improved efficiency of iPSC generation 4-to 10-fold in various reprogramming contexts. We revealed that ZFP266 binds to B1 SINEs adjacent to OSK binding sites during reprogramming and impedes chromatin opening. Furthermore, replacing its KRAB co-suppressor interacting domain with a co-activator interacting domain converted ZFP266 from a reprogramming inhibitor to a reprogramming facilitator. This indicated that B1 SINEs next to OSK binding sites during reprogramming were critical genetic elements that modulate the efficiency of OSKM-mediated iPSC generation. This work serves as a resource for better understanding of reprogramming mechanisms and highlights SINEs as novel transposable elements (TEs) involved in pluripotency induction.

### A CRISPR/Cas9-mediated genome-wide KO screen identified 16 novel reprogramming roadblock genes

We have previously generated a Cas9 expressing mouse ES cell line, named Cas9 TNG MKOS, with a *Nanog-GFP-ires-Puro* reporter and a doxycycline-inducible *MKOS-ires-mOrange* polycistronic reprogramming cassette in the *Sp3* locus (Supplemental Figure S1A and S1B)^20,21^. Efficient KO by lentiviral sgRNA delivery in both Cas9 TNG MKOS ESCs and mouse embryonic fibroblasts (MEFs), generated through morula aggregation of these ESCs, was confirmed using sgRNAs against ICAM1 and CD44, respectively, resulting in >80% loss of protein within 72 hours (Supplemental Figure S1C and S1D). Reprogramming of Cas9 TNG MKOS MEFs following sgRNA transduction against known roadblock genes *Trp53* and *Rb1*^3,4,6–8,22^, and essential genes *Pou5f1* and *Kdm6a*^23^, reproduced the expected reprogramming enhancement and reduction phenotypes (Supplemental Figure S1E-S1G), confirming that the CRISPR-based KO system as a powerful tool to investigate gene function in reprogramming. We then performed genome-wide KO screening using a previously published lentiviral sgRNA library^24^, with an optimized reprogramming condition consisting of 8 days of reprogramming factor expression followed by 8 days of puromycin selection for *Nanog*-GFP^+^ iPSCs (Figure 1A and Supplemental Figure S1H-S1K). This condition resulted in an average coverage of ~170 MEFs/sgRNA/screening replicate. Genomic DNA from flow-sorted *Nanog*-GFP^+^ iPSCs was then collected in triplicate, and integrated sgRNAs were Illumina-sequenced after PCR amplification alongside the original sgRNA plasmid library.

**Figure 1.**
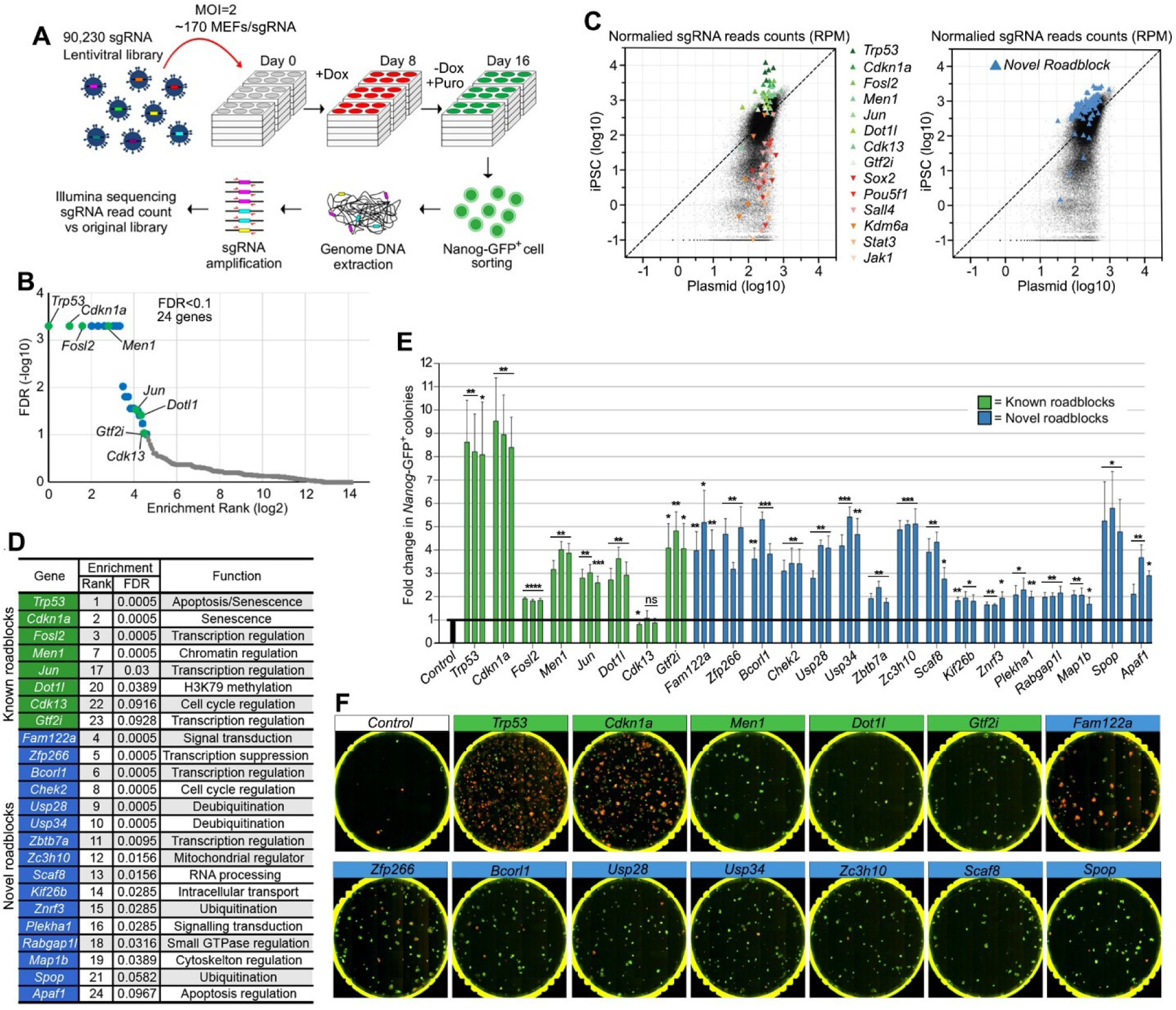
A genome-wide CRISPR screen identifies reprogramming roadblock genes. **A**. Schematic diagram of the screening strategy. sgRNA library infected Cas9 TNG MKOS MEFs were cultured in +dox for 8 days then in - dox and +Puro for additional 8 days. Integrated sgRNAs were amplified from *Nanog*-GFP^+^ cells for Illumina sequencing. **B**. Enrichment FDR ranking with MAGeCK. 24 genes, including 8 previously reported (green) and 16 novel (blue) roadblock genes, were identified using a cut-off of FDR<0.1. **C**. Normalized sgRNA read counts in the initial plasmid library versus mutant iPSC pool. sgRNAs against previously reported roadblock genes (red/orange) and genes essential for reprogramming (green) exhibited expected enrichment/depletion respectively (left). sgRNAs against 16 novel roadblock genes identified in this screen are highlighted in blue (right). **D**. Enrichment rank, FDR, and function of the 24 discovered roadblock genes. **E**. Validation of the screen result with 3 individual sgRNAs per gene. This graph is a summary of 5 data sets shown in Supplemental Figure S2A. Error bars indicate SEM, ****p<0.0001, ***p<0.001, **p<0.01,*p<0.05 based on an unpaired two-tailed t-test **F**. Representative whole-well images of KO reprogramming of 13 top roadblocks from E. Previously reported and novel roadblock genes were labelled in green and and blue, respectively. Red; mOrange, Green; *Nanog*-GFP.

The normalized read counts of all sgRNAs and analysis of the screening results with MAGeCK^25^ are available in Supplemental Tables S1, S2 and at https://kaji-crispr-screen-updated.netlify.app. Using a false discovery rate (FDR) <0.1 as a cut-off, we identified 24 genes as reprogramming roadblocks (Figure 1B). This included 16 novel candidates as well as 5 previously characterized genes: *Trp53, Cdkn1a, Dotl1* and AP-1 transcription factor components *Jun* and *Fosl2*^26,27^, and 3 genes previously uncharacterized yet identified in other screens, *Men1, Gtf2i and Cdk13*^28,29^, signifying the robustness of our screen (Figure 1C and 1D). When the top 3 ranked sgRNAs for each gene were individually tested, transduction of *Trp53* and *Cdkn1a* sgRNAs produced the largest increase in *Nanog*-GFP^+^ colony numbers (Figure 1E and 1F), although they also significantly increased the number of partially reprogrammed colonies (Figure 1F and Supplemental Figure S2A). Transduction of sgRNAs targeting all other genes, except for the previously reported roadblock *Cdk13*, enhanced *Nanog*-GFP^+^ colony formation between 2-and 6-fold (Figure 1E and Supplemental Figure S2A), verifying the inhibitory effects of the novel reprogramming roadblock genes. Expression of the validated 23 roadblock genes during reprogramming did not follow any common particular pattern and many of them exhibited consistently low expression, compared to the common housekeeping genes or the reprogramming and pluripotency marker genes^30^ (Supplemental Figure S2B). This highlights the advantage of functional screening to identify their inhibitory effects over expression profiling.

Of the 16 novel roadblock genes, KO of 8 resulted in a >4-fold enhancement similar to or better than previously reported roadblocks (Figure 1E). We therefore further characterised these 8 novel reprogramming roadblocks; *Fam122a, Zfp266, Bcorl1, Usp28, Usp34, Zc3h10, Scaf8* and *Spop* (Figure 1F, blue), alongside the previously reported roadblocks *Trp53, Cdkn1a, Men1, Dot1l* and *Gtf2i* (Figure 1F, green), as 13 top roadblocks.

### *Zfp266* KO consistently enhances and accelerates the attainment of pluripotency

Reprogramming roadblock function is influenced by multiple elements such as the stoichiometry or expression levels of OSKM, culture conditions, and starting cell types^20^. Thus, we examined the KO effects of our 13 top roadblocks in different reprogramming contexts. We first performed *piggyBac* transposon-based reprogramming with an *MKOS* or STEMCCA (*OKSM*) reprogramming cassette^31,32^ (Supplemental Figure S3A). The STEMCCA cassette expresses lower levels of KLF4 protein due to an N-terminal truncation following a 2A peptide^33^, resulting in inefficient mesenchymal-epithelial transition (MET) and a higher proportion of partially reprogrammed cells^20,34^. *piggyBac* delivery of the MKOS cassette together with each sgRNA against all the 13 roadblocks enhanced reprogramming of *Cas9 Nanog*-GFP MEFs as seen with Cas9 TNG MKOS MEFs before (Figure 2A and 2B), despite a markedly lower KO efficiency with the *piggyBac* system compared to lentiviral sgRNA delivery (Supplemental Figure S3B). In STEMCCA-mediated *piggyBac* reprogramming, KO of all roadblocks, except *Cdkn1a, Fam122a* and *Zc3h10*, increased the number of *Nanog*-GFP^+^ colonies, though *Cdkn1a* and *Fam122a* KO drastically increased *Nanog*-GFP^−^ colony numbers (Figure 2C and 2D). In particular, the KO of *Zfp266* increased numbers of *Nanog*-GFP^+^ colonies ~10-fold with almost all colonies expressing *Nanog*-GFP (Figure 2C and 2D). When *piggyBac MKOS*+sgRNA vectors were used to reprogram *Cas9* expressing neural stem cells (NSCs), sgRNAs against *Men1, Fam122a, Zfp266* and *Usp34* increased reprogramming efficiency, with KO of *Zfp266* again leading to the greatest enhancement in NANOG^+^ colony formation (~ 5-fold) (Figure 2E and 2F). When we explored reprogramming kinetics by assessing expression changes of reprogramming markers, CD44, ICAM1 and *Nanog*-GFP^35^ using Cas9 TNG MKOS MEFs, KO of 5 genes, *Men1, Gtf2i, Dot1l, Zfp266* and *Zc3h10*, demonstrated accelerated reprogramming (Figure 2G, 2H, and Supplementary Figure 3C). In summary, KO of *Zfp266* exhibited the most context-independent and robust reprogramming enhancement amongst all roadblock genes we identified. We therefore investigated further how *Zfp266* impedes the reprogramming process.

**Figure 2.**
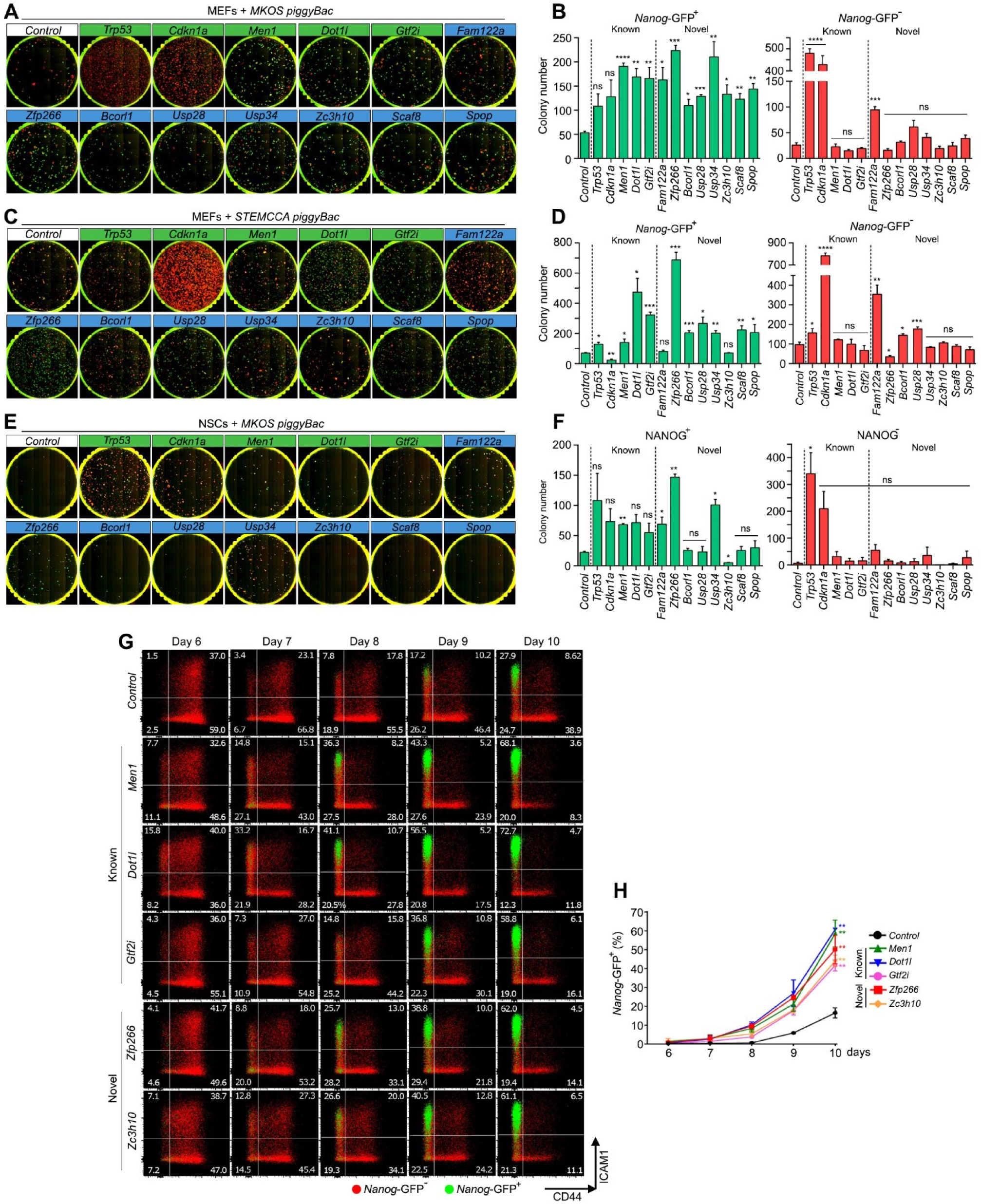
Characterization of the roadblock gene KO in different reprogramming systems and kinetics. **A, C**. *Cas9* expressing *Nanog*-GFP MEF reprogramming with MKOS (A), STEMCCA (C) *piggyBac* transposons with sgRNA expression at day 15. Red; mOrange, Green; *Nanog*-GFP. **B**,**D**. *Nanog*-GFP^+^ and *Nanog*-GFP^−^ mean colony numbers from A and C. **E**. *Cas9* expressing NSC reprogramming with MKOS *piggyBac* transposons with sgRNA expression at day 15. Green; immunofluorescence for NANOG. **F**. NANOG+ and NANOG-mean colony numbers from E. **G**. Accelerated CD44/ICAM/*Nanog*-GFP expression changes by sgRNA expression against the roadblock genes (n=2). Red; *Nanog*-GFP-cells, Green; *Nanog*-GFP+ cells. **H**. Quantification of *Nanog*-GFP+ cells from day 6 to 10 of reprogramming. The graph represents an average of 2 independent experiments. For all graphs error bars indicate SEM, ****p<0.0001, ***p<0.001, **p<0.01,*p<0.05 based on an unpaired two tailed t test.

**Figure 3.**
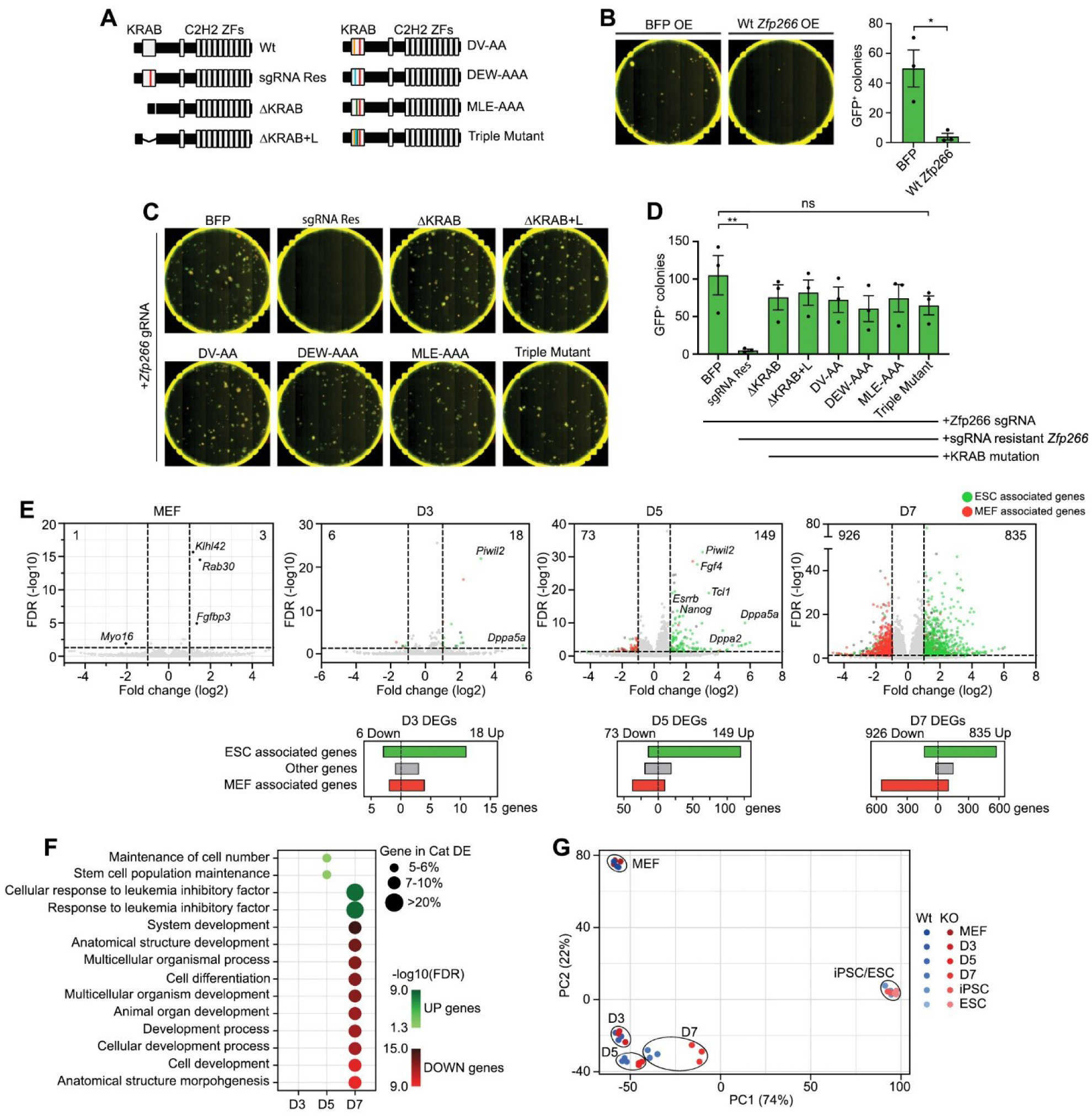
ZFP266 impedes activation of pluripotency genes via its KRAB domain. **A**. Diagram of *Zfp266* Wt and mutants. A red bar indicates a silent mutation that confers sgRNA resistance. KRAB domain deletion mutants with (Δ KRAB+L) and without a linker (Δ KRAB) do not have the sgRNA target sequence. DV-AA, DEW-AAA, MLE-AAA mutants have alanine substitutions in the indicated critical amino acids in the KRAB domain. Triple mutant contains all the alanine substitutions. **B**. *Nanog*-GFP MEF reprogramming with MKOS *piggyBac* transposons and BFP or Wt *Zfp266* cDNA overexpression, imaged at day 15. Red; mOrange, Green; *Nanog*-GFP. Error bars indicate SEM, *p<0.05 based on an unpaired two-tailed t-test. **C**. *Cas9 Nanog*-GFP MEF reprogramming with MKOS *piggyBac* transposons, *Zfp266* sgRNA expression as well as BFP, Wt *Zfp266, or* mutant *Zfp266* cDNA overexpression, imaged at day 15. Red; mOrange, Green; *Nanog*-GFP. **D**. Mean *Nanog*-GFP^+^ colony numbers of C. Error bars indicate SEM, **p<0.01 based on a one-way ANOVA test. **E**. RNA-Seq volcano plot of *Zfp266* KO vs Wt MEF, day 3, 5 and 7 of reprogramming. Up-regulated and down-regulated genes in KO cells are shown to the right and left of the plot, respectively (cut-off FDR<0.05, log2FC>|1|). ESC- and MEF-associated genes (FDR<0.05, log2FC>|1| in ESCs vs MEFs) are highlighted in green and red. Graphs below volcano plots show the number of ESC-associated, MEF-associated and other genes within D3, D5, D7 reprogramming differentially expressed genes (DEGs). **F**. GO enrichment analysis of upregulated and downregulated genes in *Zfp266* KO reprogramming. **G**. Principal component analysis of *Zfp266* Wt and KO RNA-Seq samples. Blue dots indicate Wt samples, red dots indicate KO samples, three samples per timepoint.

### ZFP266 impedes activation of pluripotency genes via its KRAB domain

*Zfp266* is predicted to encode a KRAB-ZF protein with a singular KRAB-A module in the N-terminus and putative DNA binding domain with 12x C2H2 type zinc finger array in the C-terminus (Figure 3A). KRAB domains are known to interact with co-suppressor KAP-1/TRIM28, a scaffold protein that can recruit epigenetic modifiers and promote the formation of heterochromatin and transcriptional repression^36^, suggesting that ZFP266 acts as a suppressor. Consistent with the reprogramming enhancement by *Zfp266* KO, overexpression of exogenous *Zfp266* completely disrupted reprogramming (Figure 3B). Exogenous overexpression of *Zfp266* mutants either lacking the entire KRAB domain or containing point mutations which disrupt the interaction with KAP-1/TRIM28^37,38^ could not abolish *Zfp266* sgRNA-mediated reprogramming enhancement even though the *Zfp266* mutants were resistant to the sgRNA (Figure 3C and 3D). This clearly demonstrates that Zfp266 inhibits reprogramming through its KRAB-domain.

To assess further the function of *Zfp266*, we examined the gene expression changes associated with its depletion (Supplemental Tables S3). RNA-seq of MEFs 4 days after *Zfp266* sgRNA transduction revealed only 4 differentially expressed genes (DEGs) (FDR<0.05, log2FC>|1|), demonstrating that loss of ZFP266 alone was not sufficient to cause drastic gene expression changes in MEFs (Figure 3E, Supplemental Tables S4). In contrast, the number of DEGs between *Zfp266* KO and wild-type cells rapidly increased during reprogramming from 24 at day 3 to 222 at day 5, and to 1761 at day 7 (Figure 3E, Supplemental Tables S5-S7). The majority of DEGs at day 3 and day 5 were upregulated (75% and 67% respectively, Figure 3E), consistent with the predicted role for *Zfp266* as a transcriptional suppressor. Enhanced up-regulation of pluripotency-associated genes *Piwil2* and *Dppa5a* at day 3, *Nanog, Esrrb, Dppa2, Tcl1, etc*. at day 5 was already detected in *Zfp266* KO (Figure 3E). Furthermore, over 60% (11/18), 80% (120/149), 69% (575/835) of up-regulated DEGs in *Zfp266* KO cells at day 3, 5, 7 of reprogramming were genes more highly expressed in ESCs compared to MEFs (FDR<0.05, log2FC>|1|) (Figure 3E, green, Supplemental Tables S8). Gene ontology (GO) enrichment analysis identified ‘stem cell population maintenance’ in day 5 and ‘response to leukaemia inhibitory factor’ in day 7 up-regulated DEGs as the most enriched terms (Figure 3F), while downregulated DEGs at day 7 were significantly enriched in developmental and differentiation terms (Figure 3F). Principal component analysis (PCA) also indicated that gene expression changes that have occurred in *Zfp266* KO reprogramming at day 5 and 7 reflected an accelerated transition towards a pluripotent state (Figure 3G). Taken together, these data indicate that *Zfp266* KO enhances and accelerates reprogramming by permitting a more efficient activation of pluripotency genes by OSKM.

### *Zfp266* KO in MEFs results in chromatin opening at the SINE-containing Zfp266 binding sites

One possible mechanism by which ZFP266 impedes reprogramming is that ZFP266 binds and suppresses the pluripotency loci in MEFs and other differentiated cells. To investigate this possibility, we mapped ZFP266 binding sites in MEFs using DamID-seq, which does not required specific antibodies^39,40^. This identified 15,119 unique ZFP266 binding sites (Figure 4A), predominantly situated in introns or intergenic regions (Supplementary Figure 4A, Supplemental Tables S9). These ZFP266 binding sites have low chromatin accessibility as measured by ATAC-seq in MEFs, 72 hours after reprogramming, as well as in iPSCs^41^ (Figure 4A). ZFP266 binding sites were predominantly enriched for somatic AP-1 TF motifs (Figure 4B), and little OSKM binding was observed at the same loci in 48hr reprogramming or ESC ChIP-Seq datasets^42,43^ (Supplementary Figure 4B and 4C). This disputed the idea that ZFP266 functions as a suppressor at the pluripotency-related gene loci in differentiated cells. Thus, we investigated whether any TE families were enriched in ZFP266 binding sites, since many KRAB-ZFPs are known to bind and supress transcription of TEs^15^. In line with this, we found the 15,119 ZFP266 DamID-seq peaks in MEFs were highly enriched in SINEs with about two thirds (10,523) overlapping with SINEs (Figure 4C and 4D). Of SINE subfamilies, B1 SINEs in particular exhibited both the most significant enrichment and the most abundant overlap with ZFP266 binding sites (Figure 4C and 4D). Furthermore, de novo motif analysis of ZFP266 binding sites identified 3 long de novo motifs which all corresponded to parts of the B1 SINE consensus sequence (Figure 4E), suggesting ZFP266 might bind B1 SINEs.

**Figure 4.**
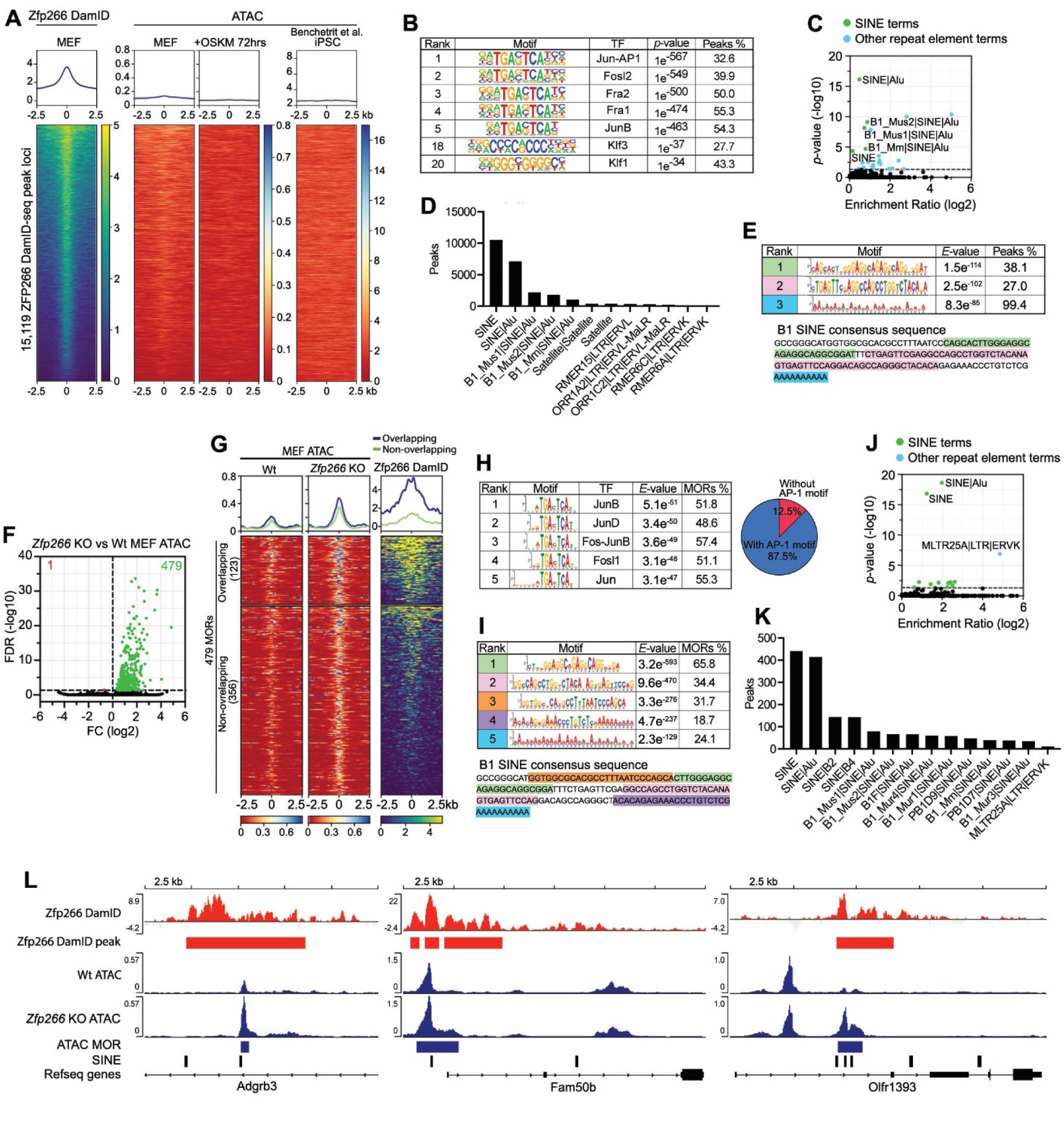
*Zfp266* KO in MEFs results in chromatin opening at the SINE-containing ZFP266 binding sites. **A**. ZFP266 DamID-seq signals in MEFs, ATAC-seq signals in MEFs, +72hours of reprogramming, and iPSCs at the ZFP266 DamID-seq peak loci. **B**. Motif enrichment analysis with HOMER on ZFP266 DamID-seq peaks. **C**. Significance and fold enrichment ratio of transposable element (TE) families overlap with Dam-ZFP266 peaks. Green dots indicate significantly enriched SINEs, blue dots indicate other significantly enriched TEs. **D**. Number of ZFP266 DamID-seq peaks that overlap with TEs. **E**. *De novo* motif discovery analysis with MEME on ZFP266 DamID-seq peaks. The identified motifs correspond to parts of the B1 SINE consensus sequence, indicated by matching colours. **F**. Volcano plot of *Zfp266* KO vs Wt MEF ATAC-seq. Green and red dots indicate more open regions (MORs) and more closed regions in *Zfp266* KO MEFs, respectively (FDR<0.05). **G**. ATAC-seq and ZFP266 DamID-seq signals in the *Zfp266* KO MEF MORs, overlapped (top) and non-overlapped (bottom) with ZFP266 DamID peaks in MEFs. **H**. Motif enrichment analysis on *Zfp266* KO MEF MORs. Percentages of MORs containing each motif and AP-1 motif are indicated. **I**. *De novo* motif discovery analysis on *Zfp266* KO MEF MORs. The top five most significant motifs correspond to parts of the B1 SINE consensus sequence, indicated by matching colours. **J**. Significance and fold enrichment ratio of transposable element (TE) families overlap with Zfp266 KO MEF MORs. Green dots indicate significantly enriched SINEs, blue dots indicate other significantly enriched TEs. **K**. Number of *Zfp266* KO MEF MORs that overlap with TEs. **L**. Examples of *Zfp266* KO MEF MORs (Blue) with SINE (black), overlapping with ZFP266 DamID-seq peaks (red).

We next examined how depletion of ZFP266 might affect chromatin accessibility. To this end, we performed ATAC-Seq of *Zfp266* KO MEFs and identified 479 more open regions (MORs) compared to WT MEFs, while only one locus was found to be a more closed (Figure 4F and Supplemental Tables S10). Considering the predicted suppressor function of ZFP266, we next examined whether ZFP266 binds to the MORs in wild-type MEFs. Although only about 25% (123/479) of *Zfp266* KO MEF MORs overlapped with ZFP266 DamID-seq peaks, non-overlapped MORs also had increased DamID-seq signals albeit at a lower level (Figure 4G), unlike randomly selected control regions with similar chromatin accessibility (Supplementary Figure 4D). This suggests that more than 25% of MORs are likely bound by ZFP266, while they were not identified as a ‘peak’ with our DamID-seq due to the cut-off criteria and/or technical limitations. In fact, similar to ZFP266 DamID-seq peaks in MEFs, *Zfp266* KO MEF MORs were mainly located in intergenic regions and introns (Supplementary Figure 4E), and enriched in AP-1 TF motifs, with 87% of all *Zfp266* KO MEF MORs containing at least one AP-1 TF motif (Figure 4H). *De novo* motif discovery analysis also identified 5 motifs that overlap with the B1 SINE consensus sequence (Figure 4I), consistent with the fact that SINEs were the most enriched repetitive element (Figure 4J), and 92% (441/479) of *Zfp266* KO MEF MORs had at least one SINE (Figure 4K). Overall, these data suggest that ZFP266 binds to B1 SINEs in MEFs to keep target loci closed, and removal of ZFP266 allows TFs that binds nearby, like AP-1 TFs, to facilitate chromatin opening (Figure 4L). However, ZFP266 does neither bind to nor regulate pluripotency gene loci in MEFs.

### Zfp266 KO in reprogramming results in chromatin opening at SINE-containing OSK binding sites

In order to address why *Zfp266* KO results in significant reprogramming enhancement, we performed ATAC-seq 72 hours after reprogramming with and without *Zfp266* KO. Similar to the KO effects in MEF, *Zfp266* KO reprogramming cells exhibited 1522 MORs, the majority of which were situated in intergenic regions and introns (Supplementary Figure 5A), and only 86 more closed regions compared to wild-type cells (Figure 5A, Supplemental Tables S11). They were also significantly enriched in SINEs, particularly B1 SINEs (Figure 5B), with >90% (1459/1522) of MORs containing at least one SINE (Figure 5C). *De novo* motif discovery analysis also identified motifs that correspond to the B1 SINE consensus sequence as the most significant motifs (Figure 5D). However, these loci hardly overlapped with *Zfp266* KO MEF MORs (Figure 5E), suggesting a context dependency for which loci become more open in the absence of ZFP266. The overlap with MEF ZFP266 DamID-seq peaks was also minimal, with only ~10% of MORs overlapping (Supplemental Figure S5B), and non-overlapped MORs did not have increased DamID-seq signals compared to the control regions with a similar chromatin accessibility (Supplemental Figure S5B). This indicated that upon OSKM expression ZFP266 changes binding sites at which it regulates chromatin accessibility. TF motif enrichment analysis revealed that KLF, SOX and the OCT4::SOX2 motifs were highly enriched in *Zfp266* KO reprogramming MORs, particularly with KLF family (KLF1, KLF5, KLF4, KLF9, KLF12) motifs identified in >90% of these MORs, while AP-1 TF motifs were also enriched (Figure 5F). Next, we classified *Zfp266* KO reprogramming MORs into two groups using *K*-means clustering (Figure 5G). The first cluster (121 regions) are open in both wild-type and *Zfp266* KO MEFs, and then become more closed upon reprogramming, while *Zfp266* KO cells are more resistant to this closing (cluster 1, Figure 5G). The second cluster contained the majority (1401) of *Zfp266* KO reprogramming MORs, which are closed in both wild-type and *Zfp266* KO MEFs, and become open following reprogramming, an effect which is enhanced when *Zfp266* is knocked out (cluster 2, Figure 5G). Cluster 2 indicates that removal of ZFP266 facilitates OSKM-mediated chromatin opening. In fact, we observed OSK binding in cluster 2 reprogramming MORs with particularly strong KLF4 signal using published reprogramming 48-hour ChIP-Seq datasets^43^ (Figure 5H). Similar OSK binding was observed in an ESC ChIP-Seq dataset albeit to a lesser extent^42^ (Supplemental Figure S5C), and about one third of the cluster 2 loci have an open chromatin state in iPSCs (Supplemental Figure S5D), suggesting that some of the MORs are OSK targets in pluripotent cells. Interestingly, while both OSK binding and motifs were enriched close to the MOR peak summit (within 70 bp) (Figure 5H and 5I), SINEs were depleted from summits and were instead enriched immediately upstream or downstream (~70 bp away from the summit), therefore being located immediately adjacent to OSK motifs and binding sites (Figure 5I and 5J).

**Figure 5.**
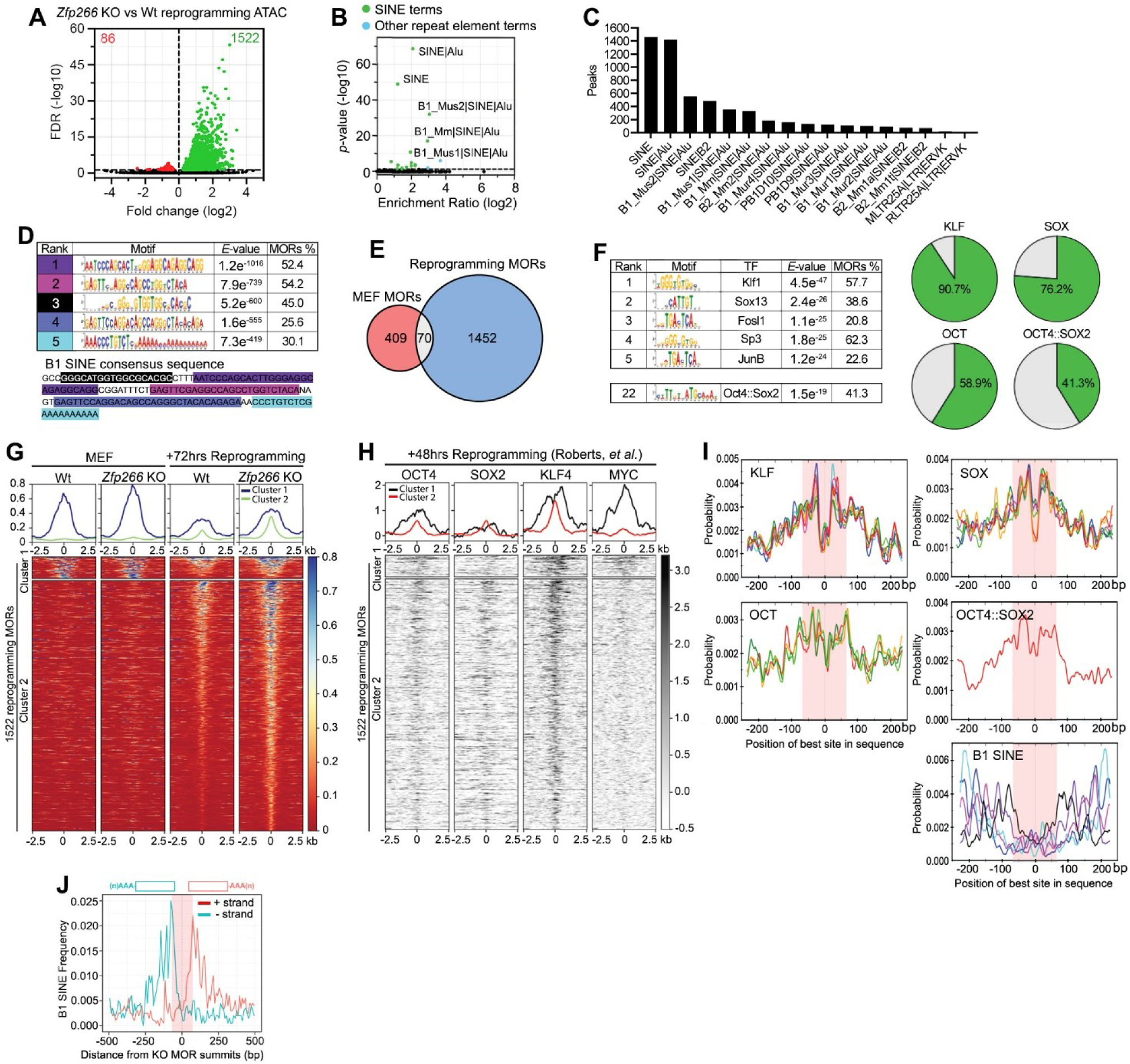
Reprogramming *Zfp266* KO MEFs results in chromatin opening at OSK bound, B1 SINE containing loci. **A**. ATAC-seq volcano plot of *Zfp266* KO vs Wt reprogramming (+72hours after OSKM induction). Green and red dots indicate more open regions (MORs) and more closed regions in *Zfp266* KO reprogramming, respectively (FDR<0.05). **B**. Significance and fold enrichment ratio of transposable element (TE) families overlap with Zfp266 KO MEF MORs. Green dots indicate significantly enriched SINEs, blue dots indicate other significantly enriched TEs. **C**. Number of *Zfp266* KO reprogramming MORs that overlap with TEs. **D**. *De novo* motif discovery analysis on *Zfp266* KO reprogramming MORs. The motifs correspond to parts of the B1 SINE consensus sequence are indicated by matching colours. **E**. Overlap between *Zfp266* KO MEF MORs and *Zfp266* KO reprogramming MORs. **F**. Motif enrichment analysis with *Zfp266* KO reprogramming MOR peak summits and percentages of MORs with KLF, SOX, OCT (POU) family and OCT4::SOX2 motifs. **G**. Classification of *Zfp266* KO reprogramming MORs based on ATAC-seq signals in MEF and reprogramming 72 hours. **H**. Reprogramming 48 hours OSKM ChIP-Seq heatmap plots at *Zfp266* KO reprogramming MORs. **I**. KLF, SOX, OCT (POU) family, OCT4::SOX2 and SINE motif distribution within *Zfp266* KO reprogramming MORs. 70 bp from the summit is highlighted in pink. The colours of B1 SINE motifs correlate to those in D. **J**. Orientation-biased distribution of B1 SINE elements within *Zfp266* KO reprogramming MORs. Head of B1 SINE tends to locate on the MOR summit side. 70 bp from the summit is highlighted in pink.

Furthermore, B1 SINEs within the MORs were mostly orientated such that the 5’ head sequence was positioned inwards facing towards the peak summit (Figure 5I and 5J). This positional and directional bias within the MOR was exclusive to B1 SINE subfamilies as B2 SINEs exhibited no such bias (Supplemental Figure S5E). Based on these data together with a report that somatic TFs’ binding sites drastically change upon OSKM expression^42^, we speculated that ZFP266 was relocated to B1 SINEs next to OSK binding sites during reprogramming, where it then impeded chromatin opening.

### Facilitating chromatin opening at ZFP266 targeted SINEs enhances reprogramming

In order to validate binding of ZFP266 to SINEs, we generated an activator version of *Zfp266* with the KRAB domain replaced by a flexible linker and three transactivating domains VP64, p65 and Rta (VPR)^44^ (Figure 6A), and performed luciferase reporter assays using HEK293 cells. Enhanced luciferase expression was observed when VPR-*Zfp266*, but not BFP or VPR only controls, was co-transfected with a reporter plasmid containing the B1 SINE consensus sequence upstream of a SV40 minimal promoter (Figure 6B). Co-expression of wild-type *Zfp266* alongside VPR-*Zfp266* attenuated this reporter expression (Figure 6C), confirming ZFP266 specifically binds B1 SINEs. We next examined whether VPR-ZFP266 can bind to *Zfp266* KO reprogramming MORs using the *Luciferase* assay. We selected SINE containing MORs in three genes, *B3gnt3, Piwil2* and *Snx20*, whose transient up-regulation during reprogramming was significantly augmented by *Zfp266* KO (Figure 6D). These loci are closed in MEFs, open up more in *Zfp266* KO cells upon reprogramming, and are bound by KLF4 at 48 hours of reprogramming (Figure 6E). Each MOR was cloned upstream of a minimal SV40 promoter in both a forward and reverse orientation in a luciferase reporter vector. We found that VPR-*Zfp266* could also enhance luciferase expression from these MOR reporter vectors (Figure 6F), while co-expression of wild-type *Zfp266* would then ablate it (Figure 6G). Deleting B1 SINE sequences from the *B3gnt3, Piwil2* and *Snx20* MORs diminished VPR-*Zfp266*’s ability to enhance *luciferase* expression, confirming that ZFP266 binds to reprogramming MORs specifically via B1 SINE sequences (Figure 6H-6J). We also confirmed that OSKM expression enhanced luciferase expression from the *B3gnt3* and *Snx20* MOR containing reporter vectors in MEFs (Figure 6K), indicating the MORs have OSKM-dependent enhancer activity. In addition, removing B1 SINE sequences from the *Snx20* MOR led to an increase in *Luciferase* expression following OSKM induction (Figure 6L), demonstrating B1 SINEs function to repress reprogramming factor-mediated transactivation, presumably via endogenous ZFP266 binding. Finally, overexpression of the VPR-*Zfp266* together with OSKM lead to accelerated and enhanced reprogramming with a robust appearance of *Nanog*-GFP^+^ colonies by day 9 (Figure 6M and 6N). Taken together, we propose a model where 1) ZFP266 binds to B1 SINEs adjacent to OSK binding sites during reprogramming and acts to impede chromatin opening, and 2) KO of *Zfp266* (or recruitment of co-activators to these loci) tips the balance in favour of OSK, allowing them to establish a more open chromatin state to drive gene activation necessary for successful reprogramming (Figure 6O).

**Figure 6.**
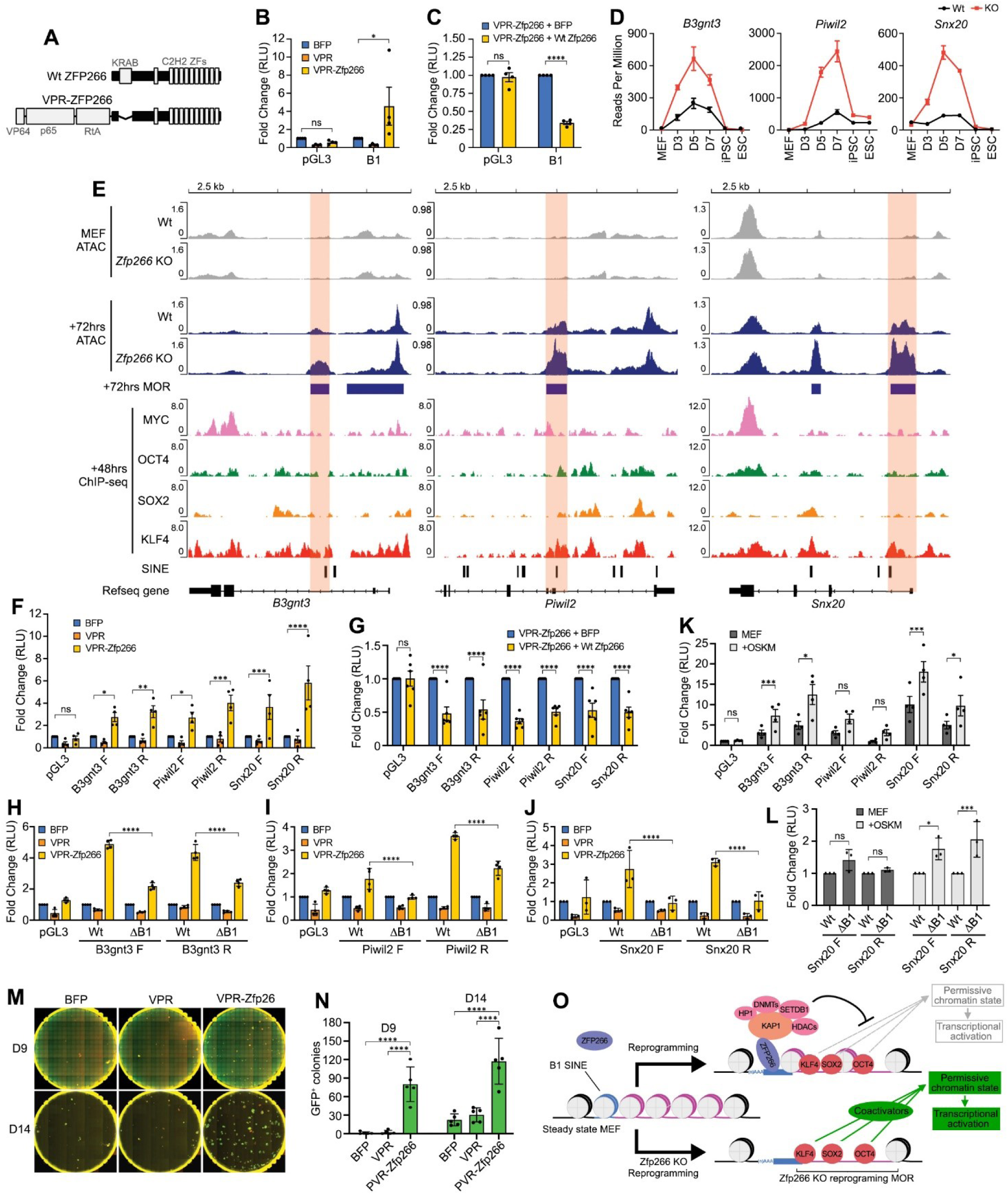
ZFP266 binds to B1 SINEs in Zfp266 KO reprogramming MORs that impede OSKM-mediated gene activation. **A**. Diagram of Wt ZFP266 and a synthetic activator version of ZFP266, VPR-ZFP266. **B, C**. Luciferase reporter assay with either an empty reporter vector pGL3 or with a reporter vector with the B1 SINE consensus sequence, co-expressed with either BFP, VPR only or VPR-*Zfp266* expression vectors (**B**), with either BFP or Wt *Zfp266* expression vectors in the presence of VPR-ZFP266 (**C**) in HEK293 cells. RLU: Relative Light Units, *p<0.05, ****p<0.0001 based on a two-way ANOVA test. **D**. *B3gnt3, Piwil2* and *Snx20* mRNA expression from the *Zfp266* Wt and KO reprogramming RNA-seq data. **E**. ATAC-seq, ChIP-seq signals at the *B3gnt3, Piwil2* and *Snx20* MORs, cloned in both forward (F) and reverse (R) directions (relative to gene orientation) for luciferase reporter assays (highlighted in orange). **F-J**. Luciferase reporter assay with an empty reporter vector pGL3 or vectors containing *B3gnt3, Piwil2* and *Snx20* MORs co-transfected with either BFP, VPR only or VPR-*Zfp266* expression vectors (**F**), co-transfected with either BFP or Wt *Zfp266* expression vectors in the presence of VPR-ZFP266 (**G**), an empty reporter vector pGL3, vectors containing *B3gnt3* (**H**), *Piwil2* (**I**) and *Snx20* (**J**) MORs with (Δ B1) or without (Wt) B1 SINE deletion co-transfected with either BFP, VPR only or VPR-*Zfp266* expression vectors in HEK293 cells. **K, L**. Luciferase reporter assay with empty reporter vector pGL3 or vectors containing *B3gnt3, Piwil2* and *Snx20* MORs (**K**), *Snx20* MOR reporter with (Δ B1) or without (Wt) B1 SINE deletion (**L**), using MEF with or without OSKM expression. *p<0.05, **p<0.01, ***p<0.001, ****p<0.0001 based on a two-way ANOVA test. **M**. Day 9 and day 14 after OSKM induction with overexpression of either BFP, *VPR* only or *VPR-Zfp266*. Red; mOrange, Green; *Nanog*-GFP. **N**. Quantification of *Nanog*-GFP^+^ colony numbers at day 9 and day 14. ****p<0.0001 based on a one-way ANOVA test. **O**. Mechanistic model of how *Zfp266* KO enhances reprogramming. ZFP266 recruited by OSK to their target loci binds to adjacent B1 SINE and impedes chromatin opening (top). *Zfp266* KO results in increased chromatin accessibility in those loci, facilitating pluripotency gene expression.

## Discussion

Reprogramming towards iPSCs is a conflict between OSKM transcription factors trying to establish a pluripotent state and somatic factors trying to resist this disruption in cell identity. We have identified 16 novel reprogramming roadblock genes whose depletion facilitates OSKM-mediated pluripotency induction. One of the most robust roadblock genes, *Zfp266*, is recruited to OSK binding sites through the recognition of adjacent B1 SINEs, where it impedes chromatin opening via its KRAB domain. This probably underlays the accelerated up-regulation of pluripotency gene by OSKM in both *Zfp266* KO and VPR-*Zfp266* overexpression reprogramming. In fact, many of *Zfp266* KO MORs at 72 hours of reprogramming have an open chromatin state and bound by OSK in iPSCs/ESCs, and several of them are associated with pluripotency genes or other genes highly expressed in ESCs, including *Pou5f1, Sall4, Zfp42, Klf2, Piwil2, Fbxo15, Dnmt3l, Tet1/2* (Supplemental Tables S12). These genes were more efficiently up-regulated in *Zfp266* KO reprogramming compared to the control (Supplemental Tables S3). Loss of ZFP266 in MEFs lead to only 4 DEGs, suggesting that ZFP266 may not play a significant role in a static state but act as a safeguard against drastic changes of cellular states mediated by newly expressed TFs and/or extracellular ques, such as cytokines. During early embryo development, i.e. 2-8 cell stage, ICM and ESCs, SINEs, particularly B1 SINEs, are enriched in the open chromatin regions, but not in subsequent developmental stages^14^. The data from International Mouse Phenotyping Consortium shows that only 6.5% of pups from *Zfp266* heterozygous intercrosses are homozygous for the KO allele, presenting incomplete penetrance and possible roles of ZFP266 during development while there might be compensation mechanisms (https://www.mousephenotype.org/data/genes/MGI:1924769). Further investigation might reveal roles of ZFP266 in B1 SINE region closing and neighbouring gene regulation during embryo development. B1 SINEs within *Zfp266* KO reprogramming MORs displayed a distinct positioning bias enriched at the flanks of MOR summits, and an orientation bias with head towards the summits. A recent publication identified enrichment of B1 SINEs at the flanks of CD8+ T-cell specific enhancers^45^. Our work suggests that this positioning bias could be a much more general feature of B1-SINE linked regulatory elements and provides evidence that SINEs can affect chromatin states via KRAB-ZFPs. The head-to-tail orientation bias of B1 SINEs is an intriguing novel observation of our work. Stably positioned nucleosomes are highly enriched in SINE and LINE retrotransposons in human^46^. Thus, SINEs may have a further architectural or organisational role of chromatin at the regulatory elements, while it could also be influenced by surrounding DNA sequences and/or other proteins as about 20-25% of B1 SINE had the reversed tail-to-head orientation towards the MOR summits. KLF4 has been shown to bind primate/human specific TEs in naive human ESCs and during reprogramming^13,19^. Interestingly, the *de novo* motif identified in *Zfp266* KO reprogramming MORs, but not Zfp266 KO MEF MORs or ZFP266 DamID-seq peaks, has a full KLF4 binding motif with a base switch at the 5’ end of B1 SINE consensus sequence, in addition to a partial KLF4 motif enriched in nucleosome enriched KLF4 target sites^47^ (Supplementary Figure S5F). While B1 SINEs are restricted to rodents, KLF4 binding is enriched in the old world monkey-, ape-, and human-specific TEs, HERVH, HERVK, and SVA, in naïve human ESCs^19^. The enrichment of KLF4 binding in *Zfp266* KO reprogramming MORs with SINEs indicates conserved function of KLF4 to regulate gene expression via TE containing regions in reprogramming/pluripotency across species, which is of clear interest for further investigation. It has been reported that ~2/3 of human KRAB-ZFPs bind to TEs genome-wide, and KRAB-ZFPs supress not only TEs, but also expression of genes nearby^16,17^. Our results suggest a possibility that other KRAB-ZFPs could act as barriers in different TF-mediated cell conversions or differentiation of pluripotent cells to specific cell types, and therefore elimination of those obstacles or the use of engineered activator version of KRAB-ZFPs might realize more efficient cellular identity changes. Our CRISPR/Cas9-mediated genome-wide KO screening also identified several other novel genes whose inhibition of pluripotency induction would have never been predicted from transcriptomic analyses. Further understanding how these genes hamper OSKM-mediated reprogramming will bring us a better understanding of how to control cellular identities.

## Methods

### Cell culture

ESCs and MEFs were cultured in ESC medium and in MEF medium, respectively, as described previously^20^. Reprogramming was performed in reprogramming medium (ESC medium supplemented with 300 ng ml^−1^ of doxycycline (Sigma) and 10 µg ml^−1^ of L-ascorbic acid or 2-Phospho-L-ascorbic acid trisodium salt (Vitamin C) (Sigma). NSCs were cultured in NSC complete medium consisting of DMEM/F-12 Media, 1:1 Nutrient Mixture (Sigma), 1X N2 supplement (Thermo Fisher Scientific), 1X B27 supplement (Thermo Fisher Scientific), 8 mM glucose (Sigma), 100 U ml^−1^ Pencillin/Streptomycin (ThermoFisher Scientific), 0.001% BSA (ThermoScientific), 0.05 mM β-mercaptoethanol (Thermo Fisher Scientific), supplemented with 10 ng ml^−1^ mouse EGF (Peprotech) and 10 ng ml^−1^ human FGF2 (Peprotech).

### Plasmids

Plasmids used in this work are summarized in Supplemental Table S13. The plasmids and their sequences are available upon request.

### Generation of Cas9 TNG MKOS ESC line and Cas9 TNG MKOS MEFs

The *Rosa26* targeting vector carrying EF1α-hCas9-ires-neo cassette (Addgene #67987) was electroporated into TNG MKOS ESC line^20,24^. Correct targeting was confirmed by Southern blotting using KpnI and MscI digested genome DNA for a 5’ and 3’ probe, respectively. The 5’ and 3’ probes were generated from PCR amplicon using the following primers, 5’ forward CAAGTGCTCCATGCTGGAAGGATTG, 5’ reverse TGATTGGGGAGGATCCAGATGGAG, 3’ forward GGATTGCACGCAGGTTCTCCG 3’ reverse CGCCGCCAAGCTCTTCAGCAA and genome DNA (for 5’ probe) or the targeting vector (for 3’ probe) as a template. Cas9 TNG MKOS MEFs were isolated from E12.5 chimeric embryos generated via morula aggregation and the proportion of transgenic MEFs from each embryo was assessed measuring % of mOrange^+^ cells after exposing 1/10^th^ of the dissociated cells to Dox for 2 days as describe previously^20^.

### sgRNA screen

The sgRNA library (Addgene #67988) was prepared as describe before^24^. 9 × 10^6^ high contribution (>98% mOrange^+^ 2 days after addition of dox) TNG MKOS Cas9 MEF plated across 90 wells of 6-well culture plates were exposed to lentiviral sgRNA library at MOI=2 for 4 hrs. We used MOI=2 (infection efficiency ~86%) in order to increase coverage of the sgRNA library, presuming the scarcity of reprogramming relevant genes and the negligible probability of the same neutral sgRNAs being repeatedly present in combination with relevant sgRNAs. After viral containing media was removed, the cells were cultured in 3 ml of reprogramming medium. Medium was replaced once 3 ml a day for the first 3 days, and then twice 4 ml a day from day 4 of reprogramming. From day 8, the media was switched to ESC medium supplemented with puromycin (1 μg ml^−1^) and medium was replenished twice a day with 4 ml / well until day 16. Puromycin resistant, *Nanog*-GFP^+^ cells were then sorted using the FACS AriaII (BD Biosciences) and stored at −80° C as cell pellets before extraction of genomic DNA. Screening was performed in triplicate. Genomic DNA from 3×10^7^ sorted GFP^+^ cells was extracted using the Blood & Cell Culture DNA Maxi Kit (Qiagen). Amplification of sgRNA regions from the extracted genome and the original sgRNA plasmid library, and Illumina sequencing was performed as described before^48^. sgRNA read count data was analysed with MAGeCK (version 0.4.4)^25^ and genes with enriched and depleted sgRNAs were detected using the test command (default parameters).

### Cas9 TNG MKOS MEF reprogramming

0.25 × 10^4^ Cas9 TNG MKOS MEFs were mixed with 9.5 × 10^4^ WT MEFs (129 strain) and seeded in gelatine-coated wells of 6-well plates. Cells were transduced with sgRNA lentiviruses at an MOI of 3 with 8 µg ml^−1^ Polybrene (Merck-Millipore) for 4 hours and then reprogramming was initiated by addition of reprogramming medium. On day 14-16, whole well colony images were taken using the Celigo S Cell Cytometer (Nexcelom) and the number of *Nanog*-GFP^+^ and *Nanog*-GFP^−^ colonies were counted. The images shown for illustration were stitched using Celigo S Cell Cytometer and processed using ImageJ.

### *piggyBac* reprogramming of MEFs with sgRNA expression and/or *Zfp266* cDNAs

*Nanog*-GFP MEFs with or without Cas9 expression from the *Rosa* locus isolated from E12.5 embryos, or wild type MEFs were plated at 1.5×10^5^ cells per well in a gelatine-coated 6-well plate. 24 hrs later co-transfection of a Dox-inducible *piggyBac* transposon vector carrying the *tetO-MKOS-ires-mOrange* or *tetO-STEMCCA-ires-mOrange* cassette with sgRNA expression cassette, *PB-CA-rtTA* vector with/without carrying a *P2A*-linked *Zfp266* cDNAs, and *pCMV-hyPBase* was performed using 500 ng each DNA and 6 μl of FugeneHD (Promega) as per manufacturer’s instructions^20,49,50^. 24 hrs later reprogramming was initiated with reprogramming medium. Medium was changed every 2 days. For colony counting, whole well colony images were taken on day 14-16 using the Celigo S Cell Cytometer (Nexcelom) and colonies were counted with ImageJ.

### *piggyBac* reprogramming of NSCs with sgRNA expression

A GFP sgRNA vector was delivered into Cas9 and GFP expressing NSCs using nucleofection with the SG Cell Line 4DNucleofector X Kit (Lonza) as per manufacturer’s instructions^51^. GFP^−^ NSCs were sorted using the FACS AriaII (BD Biosciences) and plated at clonal density. Individual clones were picked and genotyped to confirm GFP KO. NSCs were reprogrammed by nucleofection of a Dox-inducible *piggyBac* transposon vector carrying the *tetO-MKOS-ires-mOrange* cassette with/without a sgRNA expression cassette, *PB-CA-rtTA* vector and *pCMV-hyPBase*. 2×10^5^ NSCs for essential gene expression were nucleofected with 750 ng each of the above-mentioned plasmids using SG Cell Line 4DNucleofector X Kit (Lonza), DN-100 program, as per manufacturer’s instructions. Cells were recovered in NSC medium and then plated on a layer of wild type MEF feeder cells seeded the day before at a density of 1 ×10^5^ cells per well in a gelatin-coated 6-well plate. One day post-nucleofection, reprogramming was initiated with NSC complete medium supplemented with 100 U ml^−1^ human LIF, 0.3µg ml^−1^ of doxycycline (Sigma) and 10 µg ml^−1^ of L-ascorbic acid or 2-Phospho-L-ascorbic acid trisodium salt (Sigma) (sigma). After 6 days, the medium was switched to serum-free N2B27-based medium (containing DMEM/F12 medium supplemented with N2 combined 1:1 with Neurobasal® medium supplemented with B27; all from Thermo Fisher Scientific), MEK inhibitor (PD0325901, 0.8 μM, Axon Medchem), GSK3b inhibitor (CHIR99021, 3.3 μM, Axon Medchem), 1 µg ml^−1^ of doxycycline (Sigma) and 10 µg ml^−1^ of L-ascorbic acid or 2-Phospho-L-ascorbic acid trisodium salt (sigma). At day 16 of reprogramming, immunofluorescence for NANOG was performed as follows: cells fixed with 4% paraformaldehyde for 10 minutes on day 14 were permeabilized in 0.1% Triton-X in PBS for 1 hours, blocked in 5% BSA in PBS with 0.1% Tween20 for 1 hour at room temperature, and then stained in blocking solution with a primary antibody for NANOG (eBioMLC-51, Thermofisher Scientific) overnight at 4 °C. The next day, an AlexaFluor488 conjugated secondary antibody (A-21208, Invitrogen) was applied in blocking solution for 45 minutes at room temperature before washing and imaging. Whole well images were taken using the Celigo S Cell Cytometer (Nexcelom) and colonies were counted with ImageJ.

### CD44, ICAM1, *Nanog*-GFP expression analysis during reprogramming

Cells harvested at different time points of reprogramming were stained in FACS buffer for 30 min at 4°C and washed with FACS buffer prior to acquisition with LSR Fortessa (BD Biosciences) cytometer. The following antibodies from eBioscience were used: ICAM1-biotin (Clone: 13-0541; Dilution: 1/100), CD44-APC (Clone: 17-0441; Dilution 1/300), streptavidin-PE-Cy7 (Clone: 25-4317-82; Dilution: 1/1500). Dead cells were excluded using LIVE/DEAD™ Fixable Near-IR Dead Cell Stain Kit (ThermoFisher Scientific, Dilution: 1/1500). Data were analysed using Flowjo v10.

### RNA-Seq

#### Sample Preparation

For Wt and *Zfp266* KO MEF samples, 1 × 10^5^ Cas9 TNG MKOS MEFs were transduced with either a non-targeting control sgRNA or *Zfp266* sgRNA lentivirus at an MOI of 3 with 8 µg ml^−1^ polybrene (Merck-Millipore) for 4 hours. After additional 96 hours culture in MEF media, the cells were harvested for RNA extraction. For reprogramming samples, 0.25 × 10^4^ Cas9 TNG MKOS MEFs were mixed with 9.5 × 10^4^ WT MEFs (129 strain) and seeded in gelatine-coated wells of 6-well plates. Cells were transduced with either a non-targeting control sgRNA or *Zfp266* sgRNA lentivirus at an MOI of 3 with 8 µg ml^−1^ Polybrene (Merck-Millipore) for 4 hours, before being recovered for 24 hours in MEF media. After 24 hours reprogramming was initiated by addition of reprogramming medium. Cells were harvested at day 3, day 5 and day 7 of reprogramming, respectively, and 1 × 10^5^ of mOrange^+^ OSKM expressing cells were sorted with the FACS AriaII (BD Biosciences) per sample. *Nanog*-GFP^+^ iPSCs were harvested at day 15, and sorted with the FACS AriaII (BD Biosciences) into 96-well plates. Sorted iPSCs were cultured in ESC medium with puromycin (1 μg ml^−1^) to select for transgene independent clones and KO of *Zfp266* was confirmed by genotyping. *Zfp266* KO ESCs were generated by transfecting Clone J ESCs with a *Zfp266* sgRNA plasmid expressing BFP. Single BFP^+^ ESCs were then sorted with the FACS AriaII (BD Biosciences) into 96-well plates 48 hours after transfection. Clones which became BFP-negative (i.e. shed the sgRNA plasmid) were selected and KO of *Zfp266* was confirmed by genotyping. 1 × 10^5^ iPSCs or ESCs were used for RNA preparation. Cells were homogenised with the QIAshredder kit (Qiagen) and total RNA was extracted from all samples using the RNeasy Plus Micro Kit (Qiagen). Libraries were prepped with the NEB Ultra II stranded mRNA Library prep kit (NEB). RNA-Seq libraries were sequenced with NextSeq, 75SE.

### Read processing

For each sequencing run, a quality control report was generated using FastQC and Illumina TruSeq adapter sequences were removed using Cutadapt^52^. Sequencing runs from the same biological sample were then concatenated and mapped to the GRCm38 reference genome using STAR^53^.

#### Differential analysis

For each biological sample, aligned sequencing reads were first assigned to genomic features (e.g., genes) using Rsubread^54^ and a count table was generated. Differential expression analysis was then performed with DESeq2^55^, and statistically significant genes (e.g., FDR < 0.05 and log2FoldChange > 1) were identified using the standard workflow. Importantly, although the data represents a control and treatment time-series experiment, we opted to combine the factors of interest into a single factor for easier comprehension. Gene ontology analysis for differentially expressed genes was performed using the goseq package^56^.

#### Downstream analysis

For exploratory analysis and visualization, a batch-corrected and regularized log matrix of expression values was used. The count table was first transformed to stabilize the variance across the mean using the rlog function from DESeq2 and then unwanted batch effects (e.g., library preparation date) were removed using the removeBatchEffect function from limma^57^.

### DamID-seq

#### Sample Preparation

1 × 10^**5**^ WT MEFs (129 strain) were nucleofected with either *PGK-mO-Dam* or *PGK-mO-Dam-Zfp266* plasmids using the P4 Primary Cell 4D-Nucleofector X Kit (Lonza). 5 replicates were performed in total. Cells were recovered in MEF media for 48 hours before 3 × 10^**4**^ **-** 1.6 × 10^**5**^ GFP+ cells per sample were sorted with the FACS AriaII (BD Biosciences). Genomic DNA was isolated with Quick-gDNA™ MicroPrep (ZymoResearch) and 32 ng genomic DNA/sample was used for DamID-seq library preparation as previously described^39^. DamID libraries were sequenced with NextSeq, 40PE.

#### Read processing

For each sequencing run, a quality control report was generated using FastQC and Illumina Nextera adapter sequences were removed using Cutadapt. Sequencing runs from the same biological sample were then concatenated and mapped to the GRCm38 reference genome using BWA^58^. Uninformative and spurious alignments were subsequently filtered using a combination of SAMtools^59^ and BEDtools^60^ commands. Specifically, reads mapped to the mitochondrial chromosome and reads mapped to blacklisted regions were filtered.

#### Peak calling

For each biological sample, aligned sequencing reads were assigned to genomic features (e.g., DpnII restriction fragments) using Rsubread and a count table was generated. Statistically significant regions of Dam-fusion protein binding (e.g., FDR < 0.05 and log2FoldChange > 1) were detected using the callPeak command from Daim^39^. For further details, please refer to the original manuscript describing the Daim software^39^. The regions were then annotated and analysed for gene and genome ontology enrichment using the annotatePeaks command from HOMER^61^.

#### Downstream analysis

Heatmaps of read coverage at Dam-fusion binding regions were produced using the computeMatrix and plotHeatmap commands from deepTools^62^. When plotting heatmaps, a total of 5 peaks identified exactly over *Zfp266* exons (chr9:20495068-20521417) were removed from the *Zfp266* DamID peak regions due to the high signal intensity caused by the *PGK-mO-Dam-Zfp266* plasmid. *De novo* motif discovery and was performed using the MEME-ChIP tool from the MEME suite (version 5.1.1)^63^. Motif enrichment analysis was performed using findMotifsGenome command from HOMER^61^ as DamID-seq’s large peak size was not optimal for the MEME-ChIP tool. Genome browser images of peak regions and read coverage were composed using the Integrative Genomics Viewer^64^.

### ATAC-seq

#### Sample Preparation

Cas9 TNG MKOS MEFs were plated and transduced in the same manner as samples prepared for RNA-Seq. After 24 hours reprogramming was initiated by addition of reprogramming medium for reprogramming samples, while MEF samples were maintained in MEF media. Cells were harvested 96 hours after sgRNA transduction (which was 72 hours after OSKM induction for reprogramming samples) and sorted with the FACS AriaII (BD Biosciences). Cells were then processed for ATAC-Sequencing according to the Omni-ATAC protocol^65^. Briefly, 5 × 10^4^ sorted MEFs or mOrange^+^ OSKM expressing cells per sample were washed with cold 1x PBS then pelleted before the supernatant was discarded. Cell pellets were then gently resuspended in 50 µl of lysis buffer (48.5 µl resuspension buffer, 0.5 µl 10% NP-40 (Sigma) 0.5 µl 10% Tween-20 (Sigma), 0.5 µl 1% Digitonin (Promega) (resuspension buffer: 500 µl 1M Tris-HCl, pH7.5 (ThermoFisher), 100 µl 5M NaCl (Sigma), 150 µl 1M MgCl_2_ (Sigma), 49.25 ml nuclease-free H_2_O) and incubated on ice for 3 minutes. Then, 1 ml of wash buffer (990 µl resuspension buffer, 10 µl Tween-20 (Sigma)) was added to the tubes before they were gently inverted and then centrifuged for 10 minutes at 500 x g, at 4 °C. Supernatants were then carefully aspirated. Nuclei pellets were then resuspended in 50 µl of transposition mix (2.5 µl Tn5 transposase, 25 µl 2x TD buffer (both Illumina), 0.5 µl 1% Digitonin (Promega), 0.5 µl 10% Tween-20 (Sigma), 16.5 µl 1x PBS, 5µl nuclease-free H_2_O) and incubated in a thermomixer at 37 °C, 1000 rpm for 30 minutes. Transposed DNA was then purified with the Zymo DNA Clean and Concentrator-5 Kit (Zymo Research) and eluted in 21 µl nuclease-free H_2_O. All purified DNA (~20 µl) was used for PCR amplification with NEBNext High Fidelity 2x MasterMix (NEB) and optimum cycle number was determined by qPCR, as per the protocol. Amplified DNA was then purified with double-sided bead purification using AMPure XP magnetic beads (Beckman Coulter). Library concentration was determined with Qubit (ThermoFisher) and fragment size/quality with TapeStation (Agilent). ATAC libraries were sequenced with NextSeq, 40PE.

#### Read processing

For each sequencing run, a quality control report was generated using FastQC and Illumina Nextera adapter sequences were removed using Cutadapt^52^. Sequencing runs from the same biological sample were then concatenated and mapped to the GRCm38 reference genome using BWA^58^. Duplicate reads caused by PCR amplification were subsequently identified using the MarkDuplicates command from Picard (https://broadinstitute.github.io/picard/). Uninformative and spurious alignments were next filtered using a combination of SAMtools^59^ and BEDtools^60^ commands. Specifically, reads mapped to the mitochondrial chromosome, reads mapped to blacklisted regions, reads marked as duplicates, and reads not properly paired (e.g., reads that aren’t FR orientation or with an insert size greater than 2 kb) were filtered.

#### Peak calling

For each biological sample, statistically significant regions of chromatin accessibility (e.g., FDR < 0.1) were detected using the callpeak command from MACS2^66^ (https://github.com/macs3-project/MACS). For downstream analyses, a consensus set of peaks was created by taking the union across all biological samples with the multiinter command from BEDtools^60^.

#### Differential analysis

For each biological sample, aligned sequencing reads were first assigned to genomic features (e.g., consensus set of peaks) using Rsubread^54^ and a count table was generated. Differential accessibility analysis was then performed with DESeq2^67^ and statistically significant peaks (e.g., FDR < 0.05 and log2FoldChange > 1) were identified using the standard workflow.

#### Downstream analysis

Heatmaps of read coverage at chromatin accessibility regions were produced using the computeMatrix and plotHeatmap commands from deepTools^62^. K-means clustering was used to partition the regions into two distinct categories of reprogramming MORs. Genome browser images of peak regions and read coverage were composed using the Integrative Genomics Viewer^64^. Peaks were annotated against mm10 with annotatePeaks.pl from the HOMER suite (version 4.11)^61^. *De novo* motif discovery and enrichment analysis of MORs were performed using the *Zfp266* KO samples’ narrowpeak summits within MORs with the MEME-ChIP tool from the MEME suite (version 5.1.1)^63^. The number of SINE elements around peaks were counted using the BEDTools window command in a window of ±500 bp from the summits of the peaks. ATAC-seq data of iPSCs were retrieved from GSE98124^41^. ChIP-seq data of ESCs and MEFs in early reprogramming at 48 hr were retrieved from GSE90895 and GSE168142, respectively^42,43^. Chip-Seq heatmaps were generated using the deepTools computeMatrix and plotHeatmap commands^62^.

### Luciferase Reporter Assays

The pGL3 reporter plasmid containing the SV40 early promoter (Promega) was used for all luciferase reporter assays along with an internal control Renilla plasmid (Promega). Luciferase activity was measured with the GloMax 96-microplate luminometer (Promega) using the Dual-Glo Luciferase Assay System (Promega). For assays performed in HEK293 cells, 0.5-1×10^4^ cells were plated per well of a 96-well plate 24 hours prior to transfection. Transfection mixes were prepared as follows; 100 ng pGL3 reporter plasmid, 0.5 ng Renilla plasmid and 100 ng overexpression plasmid (BFP/VPR/VPR-*Zfp266*/Wt *Zfp266*) were mixed in Opti-MEM I Reduced Serum Medium (Gibco) up to 100µl. Fugene HD Transfection Reagent (Promega) was then added at a ratio of 3:1 (reagent:DNA) and 5-10 µl was added to each well of cells. Luciferase activity was measured 48 hours after transfection. For assays performed in MEF/reprogramming cells, 1×10^4^ TNG MKOS MEFs were plated per well of a 96-well plate 24 hours prior to transfection, either in MEF media or ES media +dox (300 ng/ml) to induce OSKM expression. Transfection mixes were prepared as such; 1 µ g pGL3 reporter plasmid, 10 ng Renilla plasmid were mixed in Opti-MEM I Reduced Serum Medium up to 100 µl. Fugene HD Transfection Reagent was then added at a ratio of 4:1 (reagent:DNA) and 20µl was added to each well of MEFs/reprogramming cells. Luciferase activity was measured 48 hours after transfection.

## END NOTE

## Acknowledgements

We thank I. Chambers for providing TNG ESC line, F. Rossi and C. Cryer for assistance with flow cytometry, Biomed unit staff for mouse husbandry, the Wellcome Sanger Institute sequencing facility for gRNA sequencing, EMBL GeneCore for RNA-seq, ATAC-seq and DamID-seq, A. Soufi, D. O’Caroll and M.L. Huynh for comments on the manuscript. Some of the computations for this work were enabled by resources in project SNIC 2017/7-317 provided by the Swedish National Infrastructure for Computing (SNIC) at the Uppsala Multidisciplinary Center for Advanced Computational Science (UPPMAX). This work was supported by European Research Council (ERC) grants ROADTOIPS (no. 261075) and MRC senior non-clinical fellowship (MR/N008715/1) funded for K.K. We also thank the generous support from Baillie Gifford for the collaboration between CiRA and MRC CRM, from Japan Agency for Medical Research and Development (AMED) for CiRA. K.Y. was supported by the Wellcome Trust (206194). D.F.K., J.A. and M.Y. was supported by the BBSRC (EASTBIO doctoral training partnership), Principal’s Career Development scholarship from the University of Edinburgh, and Japan Society for the Promotion of Science (JSPS) Overseas Research Fellowships, respectively. V.O. and M.Bertenstam were supported by the Swedish Foundation for Strategic Research (A3 04 159p). V.O. was also supported by the Swedish Research Council (Vr 621-2008-3074).

## Author Contribution

D.F.K. designed and performed sgRNA screen, validation and characterization of the roadblock genes including *Zfp266*. M.Y., J.A., S.K. and S.R.T. contributed to the analyses of the gRNA sequencing, RNA-seq, ATAC-seq, ChIP-seq and DamID-seq data sets. M.B. and S.Z. provided technical support. M.Bertenstam and V.O. generated the screening data website. K.Y. provided the gRNA library, the Rosa26-Cas9 targeting vector, and advised on the screen. K.K. conceived the study, supervised experiment design and data interpretation, and wrote the manuscript with D.F.K.

## Author Information

The authors declare no competing financial interests. Correspondence and requests for materials should be address to K.K.

## Notes

### Competing Interest Statement

The authors have declared no competing interest.

