## Supplemental Figs and Legends for "B1 SINE-binding ZFP266 impedes reprogramming through suppression of chromatin opening mediated by pioneering factors"

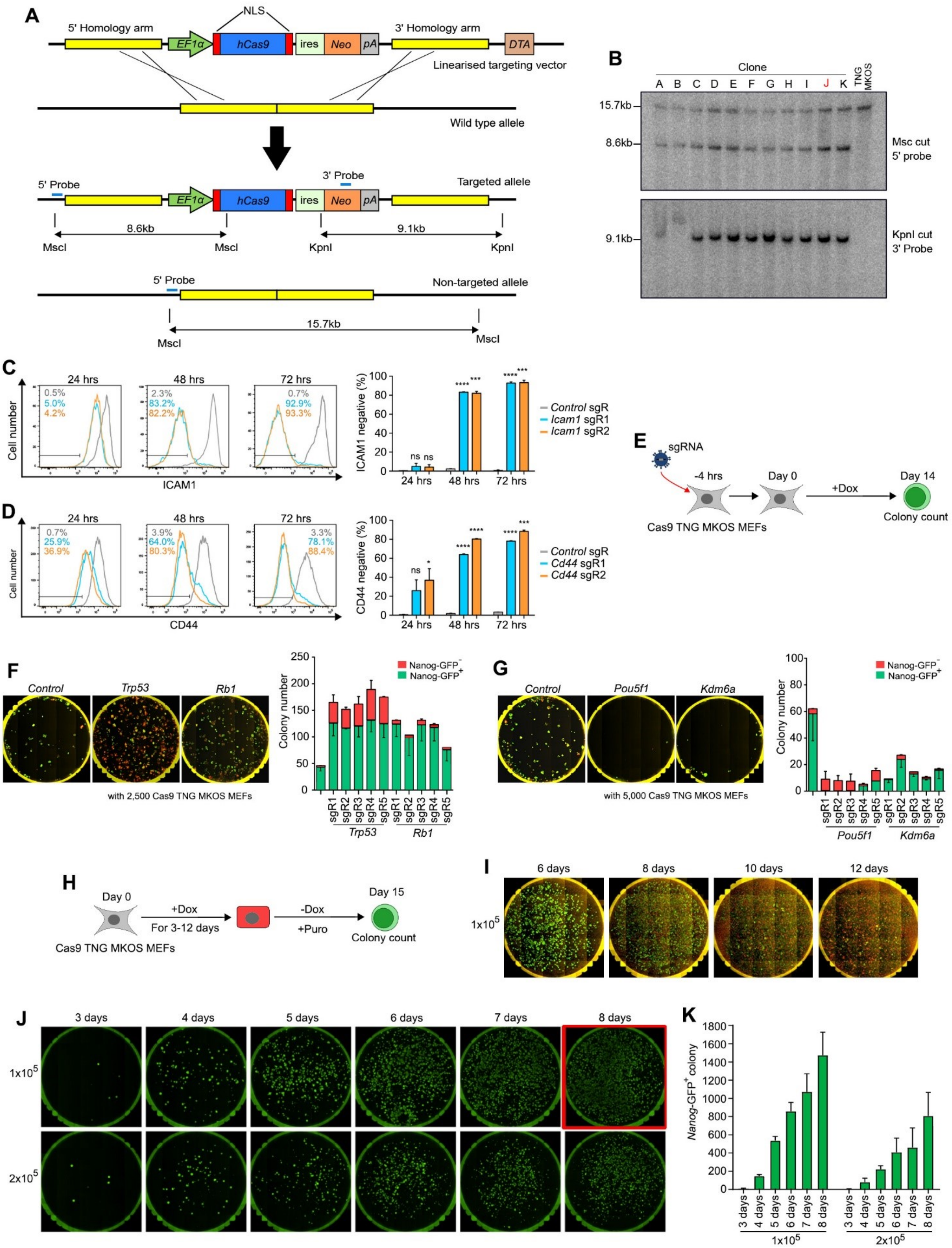


**Supplemental Figure S1. Cas9 TNG MKOS MEFs as tool for studying reprogramming. A.** Schematic diagram of the strategy to generate the Cas9 TNG MKOS ESC line. An *EF1α* promoter-driven *Cas9* expression cassette was inserted in the *Rosa26* locus of the TNG MKOS ESC line. NLS; Nuclear localisation signal, *hCas9*; Codon optimised SpCas9; ires; internal ribosome entry site, *neo*; Neomycin resistance gene, *pA;* polyadenylation signal, *DTA*; diphtheria toxin A. **B.** Southern blotting of selected G418 resistant clones. Clone J was used for all subsequent work. **C,D.** Loss of ICAM1 in TNG MKOS ESCs (**C**), and CD44 expression in Cas9 TNG MKOS MEFs (**D**), 24, 48, 72 hours after transduction of sgRNAs against *Icam1* and *Cd44*, respectively. Control sgRNA; non-targeting control sgRNA. The graph represents average of 2 independent experiments. Error bars indicate S.E.M. **p<0.01,*p<0.05, ***p<0.001, ****p<0.0001 based on a one-tailed t-test. **E.** Schematic diagram of *Cas9* TNG MKOS MEF reprogramming with sgRNA expression. Cells were cultured in +dox for 14 days, starting 4 hours after sgRNA transduction. **F.** Cas9 TNG MKOS MEF reprogramming with *Trp53, Rb1* sgRNAs. The graph represents the mean of 2 independent experiments, with 2 technical replicates. Error bars indicate SEM. **G.** Cas9 TNG MKOS MEF reprogramming with *Pou5f1, Kdm6a* sgRNA. The graph represents average of 2 independent experiments, with 2 technical replicates. Error bars indicate SEM. **H.** Schematic diagram of the optimization strategy to obtain high numbers of *Nanog*-GFP+ iPSC colonies for the screen. Cas9 TNG MKOS MEFs seeded in the absence of feeders were cultured in +dox for 3-12 days and then in -dox +Puro until day 15 for colony counting**. I.** Day 15 whole-well images of reprogramming with +dox for 6, 8, 10, 12 days starting with $1x10^5$ Cas9 TNG MKOS MEFs on day 15. Keeping +dox conditions for over 8 days decreased *Nanog*-GFP+ iPSC colony number. Red; mOrange, Green; *Nanog*-GFP. **J.** Day 15 whole-well images of reprogramming with +dox for 3 to 8 days starting with $1x10^5$, $2x10^5$ Cas9 TNG MKOS MEFs. 8 days +dox starting with $1x10^5$ *Cas9* TNG MKOS MEFs (red square) was used for the screen. **K.** Quantification of Nanog-GFP+ colonies in J. The data represents average of 2 independent experiments, with 2 technical replicates. Error bars indicate SEM.

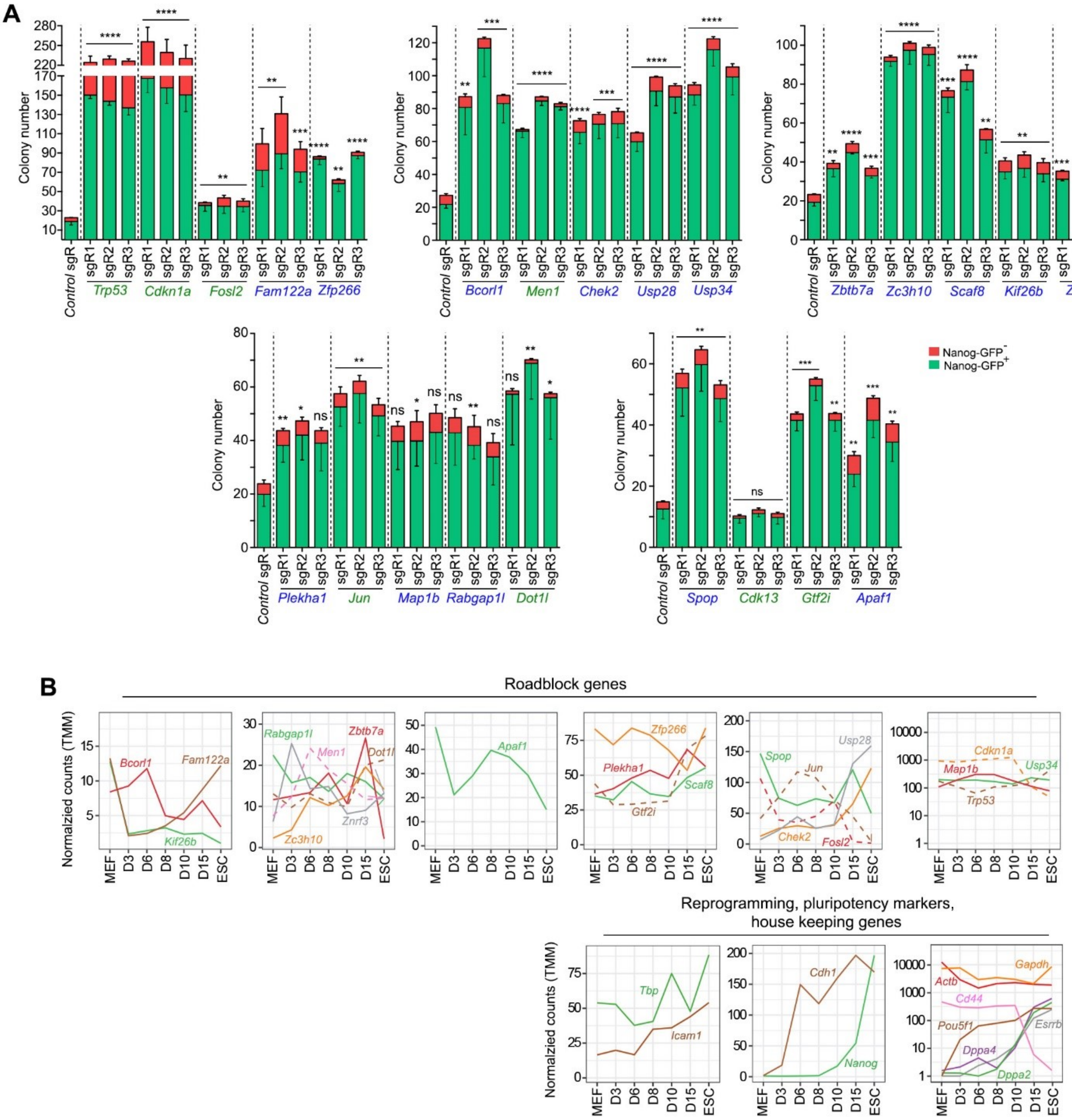


**Supplemental Figure S2. Candidate gene KO reprogramming validation and expression pattern of those genes during reprogramming. A.** Number of *Nanog*-GFP+ and *Nanog*-GFP- colonies in Cas9 TNG MKOS MEF reprogramming with sgRNA expression against the 24 roadblock genes. The data represents average of 3-4 independent experiments, with 2 technical replicates. **B.** Expression of the known (dotted line) and novel (unbroken line) roadblock genes, and reprogramming markers (*Icam1, Cdh1, Cd44*), pluripotency markers (*Nanog, Pou5f1, Esrrb, Dppa2, Dppa4*), and housekeeping genes (*Tbp, Gapdh, Actb*) during reprogramming from GEO: GSE85178[14]. The genes were grouped based on their highest expression levels.

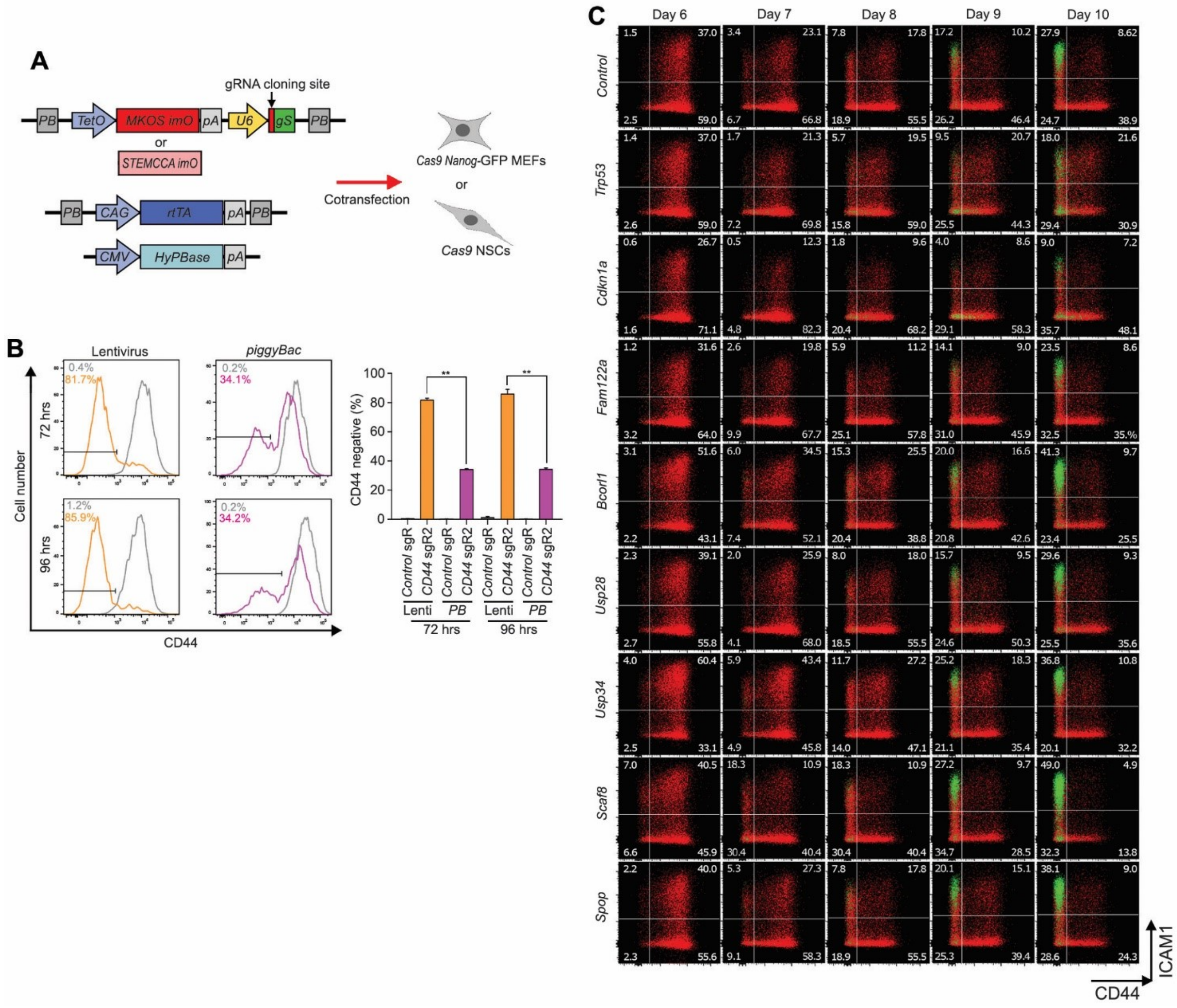


**Supplemental Figure S3. *piggyBac* reprogramming with sgRNA expression and unaffected reprogramming kinetics by KO of roadblock genes. A.** Schematic diagram of *piggyBac* reprogramming with sgRNA expression. sgRNA sequences were cloned downstream of the U6 promoter in a *piggyBac* transposon carrying the dox-inducible *MKOS-ires-mOrange* (imO) or *STEMCCA-imO* reprogramming cassette. These *piggyBac* vectors were co-transfected with a *piggyBac* vector carrying a CAG promoter-driven rtTA expression cassette and a HyPBase expression vector in Cas9 expressing *Nanog*-GFP MEFs or NSCs. gS; scaffold sequence for sgRNA. **B.** Loss of CD44 by lentiviral (left) or *piggyBac* (right) delivery of a *Cd44* sgRNA in Cas9 *Nanog*-GFP MEFs. Only cells with the sgRNA expression vector were analysed based on fluorescent reporters on the vectors. The graph represents average of 2 independent experiments. **p<0.01 based on a one-tailed t-test. **C**. CD44/ICAM/*Nanog*-GFP expression changes during reprogramming with sgRNA expression against roadblock genes (n=2). Red; *Nanog*-GFP$^-$ cells, Green; *Nanog*-GFP$^+$ cells.

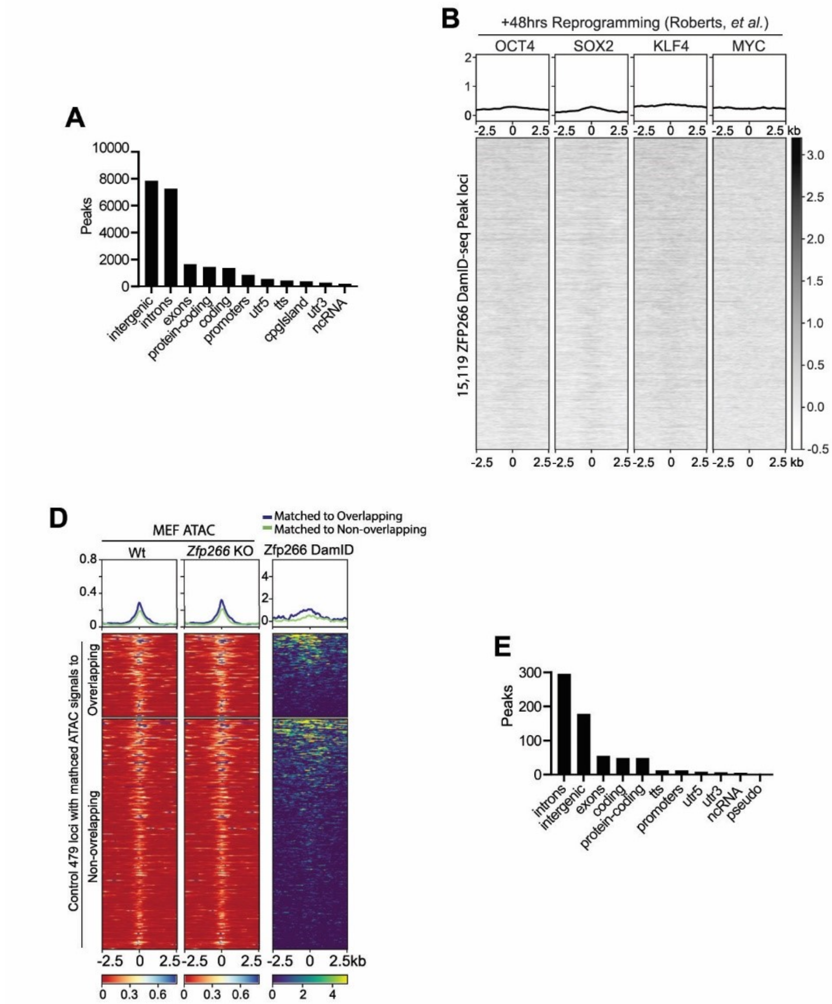


**Supplemental Figure S4. No enrichment of OSKM binding at MEF ZFP266 DamID-seq peaks and ZFP266 DamID-seq signals at *Zfp266* KO MEF MOR control regions. A.** Number of ZFP266 DamID-seq peaks that overlap with regions with the indicated genomic annotations **B, C.** OSKM ChIP-Seq signals at 48 hours of reprogramming (**B**) and ESCs (**C**) at MEF ZFP266 DamID-seq peaks. **D**. ZFP266 DamID-seq signals at *Zfp266* KO MEF MOR control regions that have similar ATAC-signals to MORs. No ZFP266 DamID-seq signal enrichment is observed in these control loci. **E**. Number of *Zfp266* KO MEF MORs that overlap with regions with the indicated genomic annotations.

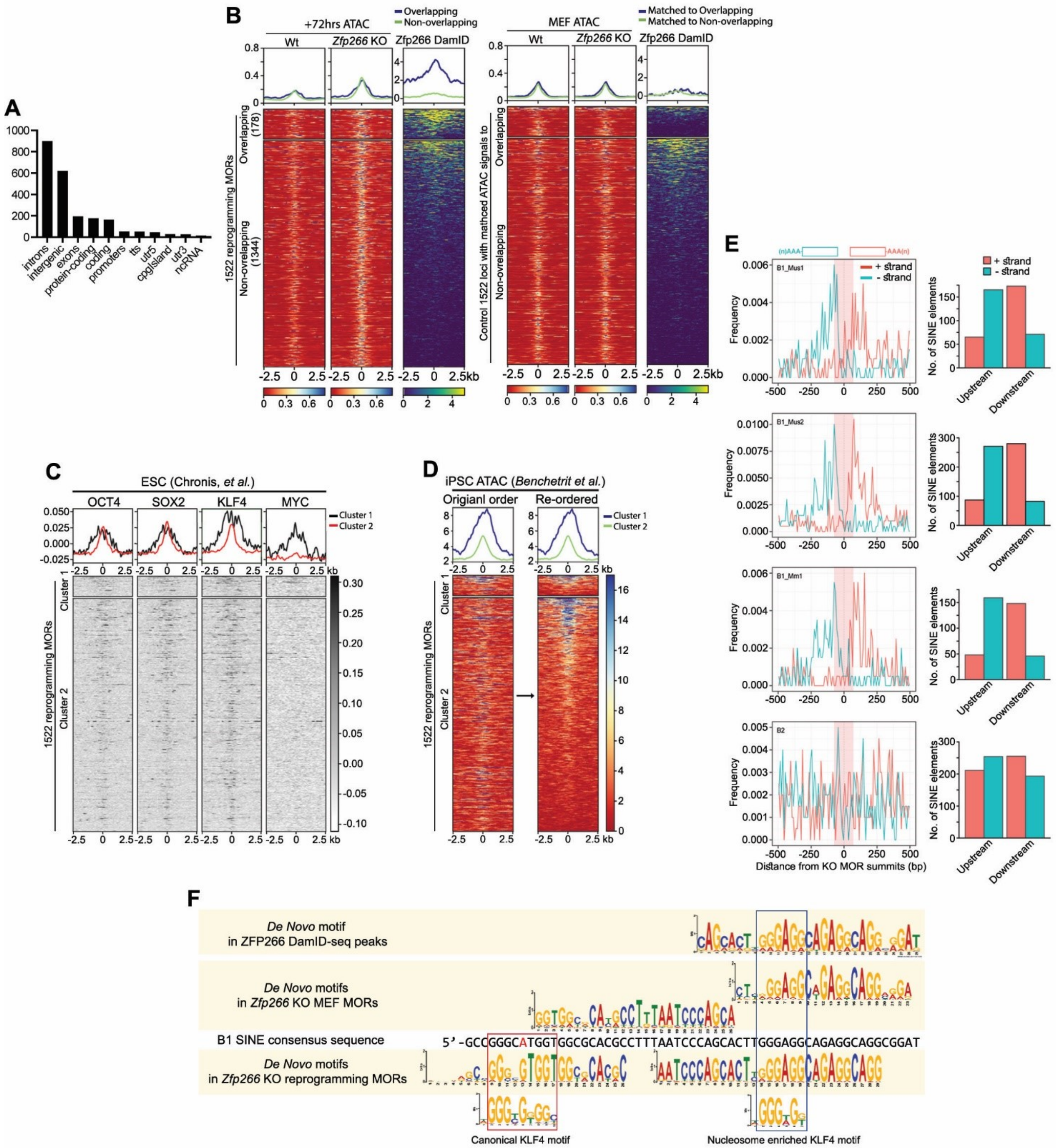


**Supplemental Figure S5. No enrichment of MEF ZFP266 DamID-seq signals, but notable OSK binding in ESCs, at majority of *Zfp266* KO reprogramming MORs. A**. Number of *Zfp266* KO reprogramming MORs that overlap with regions with the indicated genomic annotations. **B**. MEF ZFP266 DamID-seq signals at *Zfp266* KO reprogramming MORs (left) and control loci with matched chromatin accessibility (right). The DamID-seq signals at non-overlapping MORs is equivalent to those at matched control loci, unlike overlapping MORs, confirming only ~10% of reprogramming MORs are bound by ZFP266 in MEFs. **C.** OSKM ChIP-Seq signals in ESCs at *Zfp266* KO reprogramming MOR loci. **D**. ATAC-seq signals in iPSCs at *Zfp266* KO reprogramming MOR loci, in the descending order of *Zfp266* KO reprogramming ATAC-seq signals (left) or iPSC ATAC-seq signals (right). **E**. Distribution of B1 SINE subfamilies and B2 SINEs within *Zfp266* KO reprogramming MORs. Graphs on right show the actual numbers of SINEs located on the plus or minus strand either upstream or downstream of the MOR summits. **F**. Diagram showing the *de novo* motif discovered in *Zfp266* KO reprogramming MORs, but not *Zfp266* KO MEF MORs or ZFP266 DamID-seq peaks, has a canonical KLF4 binding motif with the A>G base switch at the 5' consensus sequence of B1 SINE, while a nucleosome enriched KLF4 binding motif is identified in *de novo* motifs discovered in all as well as the B1 SINE consensus sequence.
